# The exploration of *Thermococcus barophilus* lipidome reveals the widest variety of phosphoglycolipids in Thermococcales

**DOI:** 10.1101/2021.11.29.470308

**Authors:** Maxime Tourte, Sarah Coffinet, Lars Wörmer, Julius S. Lipp, Kai-Uwe Hinrichs, Philippe M. Oger

## Abstract

One of the most distinctive characteristics of Archaea is their unique lipids. While the general nature of archaeal lipids has been linked to their tolerance to extreme conditions, little is known about the diversity of lipidic structures Archaea are able to synthesize, which hinders the elucidation of the physicochemical properties of their cell membrane. In an effort to widen the known lipid repertoire of the piezophilic and hyperthermophilic model archaeon *Thermococcus barophilus*, we comprehensively characterized its intact polar lipid (IPL), core lipid (CL), and polar head group compositions using a combination of cutting-edge liquid chromatography and mass spectrometric ionization systems. We tentatively identified 82 different IPLs based on five distinct CLs and 10 polar head group derivatives of phosphatidylhexoses, including compounds reported here for the first time, e.g., di-N-acetylhexosamine phosphatidylhexose-bearing lipids. Despite having extended the knowledge on the lipidome, our results also indicate that the majority of *T. barophilus* lipids remain inaccessible to current analytical procedures and that improvements in lipid extraction and analysis are still required. This expanded yet incomplete lipidome nonetheless opens new avenues for understanding the physiology, physicochemical properties, and organization of the membrane in this archaeon as well as other Archaea.

## Introduction

Cell membranes provide dynamic physical boundaries between the inside and the outside worlds of cells of the three domains of life, Eukarya, Bacteria and Archaea. While their primary function is to ensure cellular integrity, biological membranes are much more than simple barriers: they regulate inwards and outwards fluxes, support signal transduction, cell bioenergetics and cell-to-cell communications, control cell shape, growth and division, and deform to generate, release and accept vesicles and other membrane macrostructures. These two-dimensional matrices are composite mixtures of a myriad of both lipids and proteins that are compositionally, functionally and structurally complex systems. In Eukarya, membranes are laterally organized into nano- to microscopic domains with specific compositions, physicochemical properties and functions formerly termed lipid rafts (1, 2). Such a membrane structuration is essential for membrane-hosted cellular functions, as it facilitates the organization, assembly and regulation of multimolecular protein complexes (3). Membrane order is primarily determined by membrane lipids’ tendency to phase separate (4). In Eukarya, membrane domains are thus specifically enriched in sphingolipids and cholesterol, which trigger liquid-liquid phase separation from the rest of the membrane (5, 6). However, other components and parameters have been proven essential for lateral structuration of the membrane. For instance, specific proteins such as flotillins regulate membrane domain formation (7), while the geometrical conformation of lipid polar head groups dictates their intermolecular interactions and lateral distribution (8, 9). Although they do not synthesize cholesterol, membrane lateral organization has been recently expanded to Bacteria (10), and multidimensional structuration was thus suggested to be a fundamental feature of all biological membranes (11, 12).

The membrane lipids of Archaea are however structurally divergent from those found in Bacteria and Eukarya. While the latter are typically composed of fatty-acyl ester linked to a glycerol backbone in *sn*-1,2 configuration, archaeal lipids are built upon isoprenoid cores that are ether linked to a *sn*-2,3 glycerol backbone (13–15). As a result, archaeal membranes are more stable and less permeable than those of Bacteria and Eukarya, enabling Archaea to withstand a variety of environmental conditions, ranging from the mildest to the harshest known on Earth (16, 17). Archaeal diether lipids are composed of C_15_ to C_25_ hydrocarbon chains that form bilayer membranes whereas tetraether lipids contain C_40_ side chains linked to two glycerol moieties and thus form monolayer membranes. Archaeal core lipids display a diversity of structures, which includes mono- and dialkyl glycerol diethers (MGD and DGD; (18)), glycerol mono-, di-, and trialkyl glycerol tetraethers (GMGT, GDGT, and GTGT, respectively; (19)), di- and tetraethers with hydroxylated, methylated, and unsaturated isoprenoid chains (20, 21), and tetraethers with glycerol, butanetriol, and pentanetriol backbones (22). With phosphatidic- and glycosidic polar head groups deriving from typical sugars, amino acids or combinations of both (23), archaeal polar head group diversity does not fundamentally diverge from that of Bacteria and Eukarya. However, how these diverse archaeal lipids organize into functional membranes and whether lateral organization similar to that of eukaryotic and bacterial membranes exist in Archaea remain elusive.

*Thermococcus barophilus* is a hyperthermophilic (optimal growth temperature 85 °C) and piezophilic (optimal growth pressure 40 MPa) archaeon that synthesizes both diether and tetraether lipids (24). The presence of both types of lipids implies that parts of *T. barophilus* membrane are in the form of bilayers, whereas others are monolayered, thus delineating membrane domains reminiscent of the eukaryotic and bacterial membrane lateral structuration. Additionally, the insertion of apolar polyisoprenoids in the bilayer midplane was shown to trigger lipid phase separation (25), suggesting that lateral organization is indeed possible in archaeal membranes. This model, based solely on the relative proportions of the different lipid classes in the membrane, does not account for lipid polar head groups whose charge, steric hindrance, geometry, polarity and hydrophily are critical for lipid distribution and membrane surface properties, stability, impermeability and functions (26–28). For instance, the average geometrical shape of lipids controls the propensity of these lipids to form specific phases, structures and thus domains on small to large scales (29). Resolving the exact spectrum of archaeal lipids is thus of paramount importance to grasp their biological relevance, i.e., their physiological and adaptive functions, and to comprehend membrane architecture and physicochemical properties in Archaea.

Although essential to comprehend membrane physiology, the structural diversity and distribution of archaeal lipids remain poorly characterized, partly because classic extraction procedures may lead to the preferential extraction of some classes of lipids over others (24, 30). Current data on archaeal lipids might thus not represent the real diversity in the original samples. Estimation of the lipid yield per cell indeed showed strong discrepancies with theoretical calculations of the total lipid content of different archaeal cells (31–34). For instance, intact polar lipid (IPL) extraction on *Methanothermobacter thermautotrophicus* yielded 0.038 to 0.26 fg IPL cell^-1^ whereas the theoretical lipid yield per cell for this rod-shaped archaeon (0.3 µm × 2.2 to 5.9 µm) is estimated to lie between 4.5 and 12.1 fg cell^-1^ (14). Similarly, the estimated lipid yield per cell for the coccus-shaped *Thermococcus kodakarensis* (1.1 to 1.3 µm) ranges from 7.5 to 10.8 fg cell^-1^ but IPL extraction only yielded 0.38 to 1.61 fg cell^-1^ (35). In contrast, IPL extraction on the much smaller rod-shaped archaeon *Nitrosopumilus maritimus* (0.2 µm × 0.5 to 0.9 µm) yielded similar lipid quantities than the theoretical estimation (0.9 to 1.9 fg cell^-1^ vs 0.9 to 1.5 fg cell^-1^, respectively; (32)), which suggests that archaeal lipid extraction efficiency might be impacted by physiological parameters, e.g., size and geometry of the cells, presence and characteristics of the cell envelope, as well as the lipids themselves, e.g., the nature of the polar head groups. Although cellular lipid contents were not estimated for *T. barophilus*, similar major extraction defects were also highlighted for this archaeon. The first and only characterization of the IPL signature of *T. barophilus* only reported phosphatidylinositol(PI)-DGD. In agreement with this simple IPL composition, the acid methanolysis of the total lipid extract yielded exclusively DGD (36). However, direct acid methanolysis of *T. barophilus* biomass revealed both a high proportion of tetraethers and a drastic bias of extraction and analytical procedures towards diether-based IPLs (24, 37). Most of *T. barophilus* IPLs and polar head groups thus remain uncharacterized, impeding the understanding of its membrane physiology and organization.

In an effort to solve this missing IPL enigma, we comprehensively investigated *T. barophilus* IPL and polar head group compositions and assayed its core lipid (CL) composition as a quality control of our methodology. We report the identification of up to 82 saturated and unsaturated IPLs, including the major PI-DGD, and the first characterization of several novel archaeal IPLs, notably phosphatidyl di-N- acetylhexosamine diethers and a tetraether bearing a peculiar derivative of glycosylated phosphatidylhexose. The unsuspectedly large IPL diversity of *T. barophilus* widens the Thermococcales lipid repertoire and contributes further refinements of the proposed membrane architecture in Archaea.

## Material and methods

### Microorganism and growth conditions

*Thermococcus barophilus* strain MP was isolated from the 3,550 m deep Snake Pit hydrothermal vent, on the Mid-Atlantic Ridge (36). The strain was obtained from the UBOCC (Université de Bretagne Occidentale – type Culture Collection, France). Cultures were grown under strict anaerobiosis in a rich medium established for *Thermococcales* (38), containing 3 % *w*/*v* NaCl and 10 g L^-1^ elemental sulfur, at 85°C, pH 6.8, and atmospheric pressure. The medium was reduced by adding Na_2_S (0.1% *w*/*v* final) before inoculation. Growth was monitored by counting with a Thoma cell counting chamber (depth 0.01 mm) using a light microscope (life technologies EVOS® XL Core, × 400). Under these conditions, cell concentrations of 2 × 10^8^ cells mL^-1^ were routinely achieved.

Cells of 1-L cultures in late exponential phase were recovered by centrifugation (4000 × g, 45 min, 4 °C) and rinsed twice with an isotonic saline solution (3 % *w*/*v* NaCl). A significant amount of sulfur from the growth medium was recovered alongside cells, and the cellular dried mass was thus not estimated. The cell pellets were lyophilized overnight and kept at -80 °C until lipid extraction.

### IPL extraction and UHPLC-ESI-MS analysis

IPLs were extracted using a modified Bligh and Dyer (B&D) procedure (39), as previously described (40). Briefly, dried cells were extracted with a monophasic mixture of methanol/dichloromethane/purified water (MeOH/DCM/H_2_O; 1:2.6:0.16; *v*/*v*/*v*) using a sonication probe for 15 min. After centrifugation (2500 × g, 8 min), the supernatant was collected, and the extraction procedure was repeated twice. The supernatants were pooled, dried under a N_2_ stream, solubilized in MeOH/DCM (1:5; *v*/*v*; hereafter referred to as total lipid extract; TLE), and kept at -20 °C until analysis. A significant amount of sulfur from the growth medium was extracted alongside archaeal lipids, and the total lipid dry mass was thus not estimated. IPLs separation was first performed with a hydrophilic interaction liquid chromatography (HILIC) setting. IPLs were separated on a Waters Acquity UPLC BEH Amide 1.7 µm column (150 mm×2.1 mm, Waters Corporation, Eschborn, Germany) maintained at 40 °C by ultra-high performance liquid chromatography (UHPLC) using a Dionex UltiMate 3000RS UHPLC (ThermoFisher Scientific, Bremen, Germany) instrument equipped with an auto-injector and a Chromeleon chromatography manager software following the method described by Wörmer *et al.* (41). Di- and tetraether IPLs were eluted in the same run with a flow rate of 0.4 mL min^-1^, using the following linear gradient with A [acetonitrile(ACN):DCM:formic acid(FA):ammonium hydroxide(NH3) (75:25:0.01:0.01, *v*/*v*/*v*/*v*)] and B [MeOH:H_2_O:FA:NH_3_ (50:50:0.4:0.4, *v*/*v*/*v*/*v*)]: 99 % A (2.5 min isocratic) to 95 % A in 1.5 min, then to 75 % A in 18.5 min, and finally to 60 % A in 4 min (1 min isocratic). Alternatively and when mentioned, IPLs separation was performed with a reverse phase (RP) setting. IPLs were eluted on a Waters Acquity UPLC BEH C_18_ 1.7 µm column (150 mm×2.1 mm, Waters Corporation, Eschborn, Germany) maintained at 65 °C using the following linear gradient as described by Wörmer *et al*. (41) with A [MeOH:H_2_O:FA:NH_3_ (85:15:0.04:0.1, *v*/*v*/*v*/*v*)] and B [propan-2-ol:MeOH:FA:NH_3_ (50:50:0.04:0.1, *v*/*v*/*v*/*v*)] at a flow rate of 0.4 mL min^-1^: 100 % A (2 min isocratic) to 85 % A in 0.1 min, then to 15 % A in 18 min and finally to 0 % A in 0.1 min (8 min isocratic). 100 % A was eventually held isocratically for 7 min. Samples were thawed, dried under a N_2_ stream, and dissolved in either MeOH/DCM (1:9; *v*/*v*) or MeOH/DCM (9:1; *v*/*v*) for HILIC and RP separation, respectively. The injection volume was set to 10 µL.

Detection was achieved using a maXis quadrupole time-of-flight mass spectrometer (Q-ToF-MS, Bruker Daltonics, Bremen, Germany) equipped with an electrospray ionization (ESI) source operating in positive mode. The ESI source was also operated in negative mode, although this did not provide further information on the IPL composition. Only the conditions for the MS analysis in positive mode are thus described here and were as follows: capillary voltage 4500 V, nebulizer gas pressure 0.8 bar, drying gas (N_2_) flow 4 L min^-1^ at 200 °C, mass range *m/z* 100-2000. MS calibration was performed by a tuning mixture solution (*m/z* 322.048, 622.029, 922.010, 1221.991, 1521.972, and 1821.952) introduced by loop-injection near the end of a run and an internal lock mass throughout the entire run. MS^2^ scans were automatically obtained in data-dependent mode by fragmentation of the three to ten most abundant ions at each MS scan.

Mass spectra were visualized and analyzed using the Bruker Data Analysis software by comparing the parent molecular ion masses (occurring as H^+^, NH4^+^ or Na^+^ adducts) and the characteristic fragmentation patterns with previously described ones (35, 42, 43). For quantification, 2 ng of a phosphatidylcholine C_21_-diacylglycerol (C_21_-PC) standard were added to the sample prior to injection. The response factors of bacterial mono- and digalactosyl diacylglycerols (MGDG and DGDG, respectively) relative to the injection standard C_21_-PC were used to approximate those of the detected IPLs. Calibration curves were established by injecting two times a standard solution consisting of C_21_-PC, MGDG and DGDG in six different concentrations ranging from 0.001 to 30 ng µL^-1^. Detection was achieved only at concentrations higher than 0.1 ng µL^-1^. Under our analytical conditions, MGDG and DGDG showed molecular response factors of 0.58 and 0.21 relative to C_21_-PC, respectively. Different response factors are to be expected in ESI-MS, notably for lipids bearing distinct polar head groups (e.g. hexose vs. hexosamine), but the same response factors were applied for all IPLs bearing the same number of sugar residues in their polar head group, i.e., 0.58 for all mono- and 0.21 for all diglycosylated lipids, respectively. IPL relative abundances were determined in positive mode by integration of the peak area on the extraction ion mass chromatograms with a width of 0.1 Da corresponding to the protonated, ammoniated and sodiated adducts, and subsequent correction using the corresponding response factor.

### CL extraction and UHPLC-APCI-MS analysis

In order to exhaustively analyze the CL composition of *T. barophilus*, polar head groups were removed by acid methanolysis (1 N HCl in MeOH at 70 °C for 16 h) of the biomass (total CLs) as described by Becker *et al* (44). The hydrolyzed lipids were extracted with a monophasic mixture of MeOH/DCM (1:5; *v*/*v*) using a sonication probe for 15 min. After centrifugation (2500 × g, 8 min), the supernatant was collected in a separatory funnel and the extraction procedure was repeated twice. CLs were partitioned into the organic phase following addition of Milli-Q water, and the aqueous phase was subsequently washed three times with an equal amount of DCM. The organic phases were collected, pooled, and subsequently washed three times with an equal amount of Milli-Q water. The solvent of the resulting CL extracts was evaporated under a N_2_ stream and the extracts were resolubilized in *n*-hexane/propan-2-ol (99.5:0.5; *v*/*v*). The same procedure was applied to the TLE (CLs from IPLs) to evaluate our IPL extraction method.

CLs were separated on two coupled Waters Acquity UPLC BEH Amide 1.7 µm columns (150 mm×2.1 mm, Waters Corporation, Eschborn, Germany) maintained at 50 °C using a Dionex UltiMate 3000RS UHPLC (ThermoFisher Scientific, Bremen, Germany) instrument equipped with an auto-injector and a Chromeleon chromatography manager software. The injection volume was set to 10 µL. Di- and tetraether CLs were eluted in the same run using a linear gradient with *n*-hexane and *n*-hexane/propan-2-ol (9:1; *v*/*v*) at a flow rate of 0.5 mL min^-1^, as described by Becker *et al.* (44).

Detection was achieved using a maXis Q-ToF-MS (Bruker Daltonics, Bremen, Germany) equipped with an atmospheric pressure chemical ionization (APCI) source operating in positive mode. The conditions for the MS analyses were as in Becker *et al.* (44): nebulizer gas pressure 5 bar, corona discharge current 3500 nA, drying gas (N_2_) flow 8 L min^-1^ at 160 °C, vaporizer 400 °C, mass range m/z 150-2000. MS calibration and MS^2^ scans were performed as described above.

Mass spectra were visualized and analyzed on a Bruker Data Analysis software using parent molecular ion masses (occurring exclusively as H^+^ adducts) and characteristic fragmentation patterns (45). For quantification, 2 ng of a C_46_ analogue of GTGT (C_46_-GTGT) were added to the sample prior injection. To determine the response factors of the detected core structures, calibration curves were established by injecting two times a standard solution consisting of C_46_-GTGT, GDGT with no cyclopentane ring (GDGT0) and DGD in 5 different concentrations ranging from 0.001 to 10 ng µL^-1^. GDGT0 was detected only at concentrations higher than 0.1 ng µL^-1^, whereas C_46_-GTGT and DGD were detected at all concentrations. Under our analytical conditions, DGD and GDGT0 showed relative response factors of 0.42 and 0.57 relative to C_46_-GTGT, respectively. In the absence of a measured response factor for the different archaeal core lipids, we assumed the same response factor as DGD for all diethers and that of GDGT0 for all tetraethers. CL relative abundances were determined by integration of the peak area on the extracted ion mass chromatograms with 0.1 Da width corresponding to the protonated adducts, and subsequent correction by the corresponding response factor.

### Isolation of *T. barophilus* major IPLs

In order to further characterize and validate the structural diversity of *T. barophilus* lipids, its major IPLs were isolated using the aforementioned HILIC UHPLC method. 50 % of a TLE were dried under a N_2_ stream and resolubilized in 10 µL of MeOH/DCM (1:9; *v*/*v*) for injection. Collecting vials were placed at the exit of the chromatography column, and 7 fractions corresponding to *T. barophilus* well-separated major IPLs were manually collected (F1, 11.5-12.5 min; F2, 13.3-14.0 min; F3, 14.5-15.5 min; F4, 15.7-16.4 min; F5, 17.9-18.2 min; F6, 18.4-19.0 min; F7, 23.0-27.0 min). Collected fractions were dried under a N_2_ stream, resolubilized in 100 µL of MeOH/DCM (9:1; *v*/*v*), and their purity was assayed using the aforementioned RP UHPLC-MS method. Injection volume was set to 10 µL. Detection was achieved as described in the previous section.

### IPL cleavage and UHPLC-ESI-QQQ-MS analysis of the hexose-based polar head groups

In order to characterize the hexose-based polar head groups of *T. barophilus*, both the biomass and individual IPL fractions were cleaved by acid hydrolysis (30 % trifluoroacetic acid (TFA) in H_2_O at 70 °C for 16 h) to release the monosaccharide(s) from IPLs. The reaction was stopped by drying the sample under a stream of N_2_ and the remaining TFA was removed by washing three times with DCM. Hydrolysates were solubilized in ACN/H_2_O (95:5, *v*/*v*), and CLs were extracted upon addition of hexane while monosaccharides remained in the ACN/H_2_O phase.

CLs were separated and analyzed using the aforementioned UHPLC-APCI-MS system.

Monosaccharides were separated on a Waters Acquity UPLC BEH Amide 1.7 µm column (150 mm×2.1 mm, Waters Corporation, Eschborn, Germany) maintained at 60 °C using a Dionex UltiMate 3000RS UHPLC (ThermoFisher Scientific, Bremen, Germany). Samples were dissolved in ACN/H_2_O (95:5, *v*/*v*) and the injection volume was set to 10 µL. All hexose-based polar head groups were eluted in the same run using the linear gradient described by Lowenthal *et al*. (46) with A [0.1 % NH_3_ in H_2_O] and B [0.1 % NH_3_ in ACN] at a flow rate of 0.2 mL min^-1^: 5 % A (3 min isocratic) to 10 % A in 22 min, then to 40 % A in 3 min (7 min isocratic), and finally to 5 % A in 2 min.. Detection was achieved by scheduled multiple reaction monitoring (sMRM) on a QTRAP 4500 triple quadrupole MS (ABSciEX, Darmstadt, Germany) equipped with an ESI source operating in positive mode. Source conditions were as follows: curtain gas (CUR) pressure 30 psi , ion source gas 1 (GS1) pressure 40 psi, ion source gas 2 (GS2) pressure 30 psi, ion spray (IS) voltage 4500 V, capillary temperature (TEM) 400 °C. The MRM method was established by direct infusion of 14 carbohydrates (Table S1) and consisted of 20 different transitions, with two transitions for each carbohydrate type.

The different sugar-based head groups were quantified by external calibration. Linear calibration curves were established for a wide variety of sugar derivatives by injecting two times standard solutions containing β-D-allopyranose (Alp), β-D-fructopyranose (Fru), α-D-glucopyranose (Glc), β-D-galactopyranose (Gal), α-D-mannopyranose (Man), β-D-xylofuranose (Xyl), α-L-arabinopyranose (Ara), α-D-lyxopyranose (Lyx), D-glucosamine (GlcNH_2_), N-acetyl-D-glucosamine (GlcNAc), myo-inositol (Ino) and D-saccharose (Sac) in 12 different concentrations ranging from 0.005 to 1000 µM. Alp and Fru and Xyl and Ara peaks could not be distinguished and were integrated as two single peaks (Alp/Fru and Xyl/Ara, respectively). Every standard was detected in concentration as low as 0.05 µM, with the exception of Sac which was not identified below 5 µM.

### Extraction-free analysis *via* MALDI-FT-ICR-MS

To further investigate the IPL diversity of *T. barophilus*, and particularly that of tetraether-based lipids, matrix-assisted laser desorption/ionization coupled with Fourier transform ion cyclotron resonance mass spectrometry (MALDI-FT-ICR-MS) was used directly on the biomass.

*T. barophilus* dried cell pellets were resuspended in 1 mL of Milli-Q water and centrifuged (500 × g, 1 min) to remove as much sulfur as possible. Supernatants were then decanted by centrifugation (15000 × g, 15 min) and cells were resuspended in Milli-Q water. 10 mg of 2,3-dihydroxybenzoic acid (DHB) was dissolved in 1 mL of H_2_O/ACN (3:7, *v*/*v*) containing 1 % of TFA and used as matrix. The cell suspension and the matrix solution were mixed (1:1; *v*/*v*) and 1 µL was spotted and evaporated on a ground steel MALDI target plate.

The analysis was carried out on a 7T solariX XR FT-ICR-MS coupled to a DUAL source with a Smartbeam II laser (Bruker Daltonics, Bremen, Germany). The FT-ICR-MS was operated in positive serial mode, with data being acquired over the mass range *m*/*z* 600-3000. Instrument settings were optimized for larger molecules (*m*/*z* >1000). Each scan was generated by accumulating the ions of 25 laser shots at 40 % laser power and a frequency of 2000 shots min^-1^. External calibration was performed with a standard peptide mixture (Bruker Daltonics). An internal lock mass calibration was applied using the sodiated adduct of PI- GDGT0-PI (m/z 1808.343).

## Results

### *Thermococcus barophilus* exhibits a diverse membrane lipid composition

Our B&D extracts analyzed with the UHPLC-MS procedure yielded very low cellular lipid content (0.12 fg cell^-1^, as calculated from cell counts and lipid abundance). The combined analyses of (1) the TLE and (2) the polar head groups and core structures separated from purified major lipids nevertheless revealed a diverse membrane composition for *T. barophilus* (for structures, refer to Figure 1). All IPLs identified in *T. barophilus* TLE showed ions diagnostic of archaeal phosphoglycolipids (for instance, ions at *m/z* = 733.6 and 453.3; Figure S1), whereas no glycolipids were detected.

**Figure 1:**
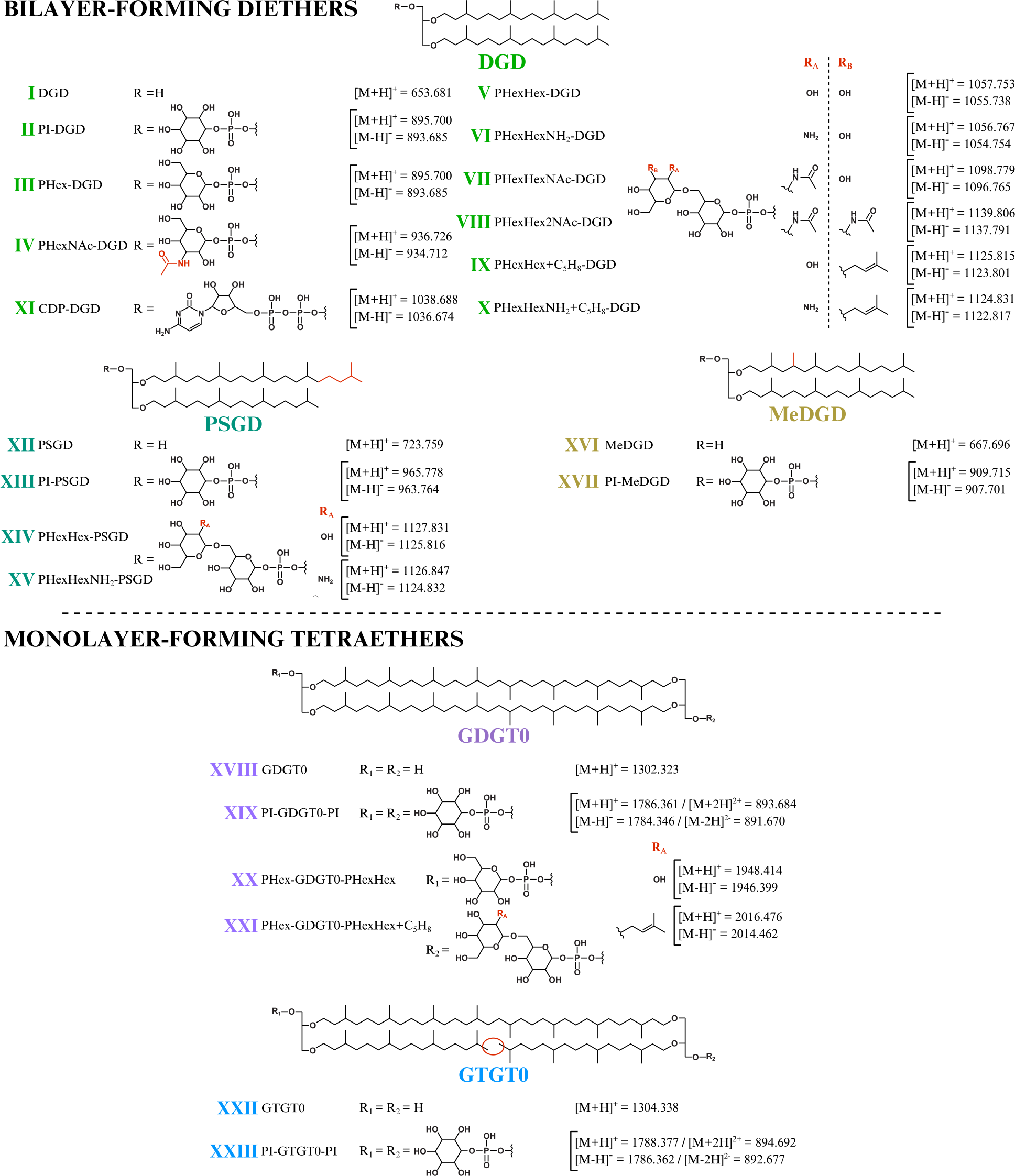
Core and intact polar lipids of *Thermococcus barophilus*. Short-hand nomenclature is indicated. The protonated, ammoniated, and sodiated adducts and only the deprotonated adducts were detected in positive and negative ion mode, respectively. Only the protonated and deprotonated ions are represented in the Figure. Core structures: diphytanyl glycerol diethers (DGD; light green; I to XI), phytanylsesterterpanyl glycerol diethers (PSGD; dark green; XII to XV), DGD bearing an additional methylation (MeDGD; yellow; XVI and XVII), glycerol dibiphytanyl (or dialkyl) glycerol tetraethers with no cyclopentane ring (GDGT0; purple; XVIII to XXI) and glycerol biphytanyl diphytanyl (or trialkyl) glycerol tetraethers with no cyclopentane ring (GTGT0; blue; XXII and XXIII). Note that II, III, IV, VI and XI were detected with up to 8 unsaturations whereas V, VII, IX, X were detected with up to 6 unsaturations. No unsaturation was detected in the other core structures. Unsaturations are not represented. Polar head groups: phosphatidylinositol (PI; II, XIII, XVII, XIX and XXIII), phosphatidylhexose (PHex; III, XX and XXI), phosphatidyl-N-acetylhexosamine (PHexNAc; IV), glycosylated phosphatidylhexose (PHexHex; V, XIV and XX), ammoniated PHexHex (PHexHexNH_2_; VI and XV), one and two N-acetylated PHexHex (PHexHexNAc and PHexHex2NAc; VII and VIII), PHexHex and PHexHexNH_2_ bearing an additional mass of 68 (PHexHex+C_5_H_8_ and PHexHexNH_2_+C_5_H_8_; IX and XXI, and X) and cytidine diphosphate (CDP; XI). Positions of additional methylation and of additional groups on the polar head groups are drawn arbitrarily in the Figure.

The TLE of *T. barophilus* appeared to be dominated by compound II (Figure 2, Table 1), whose molecular mass ([M+H]^+^ at *m/z* = 895.704) and fragmentation pattern (for instance, ions at *m/z* = 615.4 and 733.6; Figure S2) corresponded to a DGD bearing a phosphatidylhexose head group. The molecular mass and fragmentation pattern of the minor compound III (Figure S2) appeared identical to compound II but the difference in retention times (13.7 vs. 15.0 min for compounds III and II, respectively) indicated distinct hexose isomers as polar head groups of II and III. The analysis of the purified fractions revealed 100 % of Ino and 98 % of DGD for fraction F3 (98 % of compound II) and 100 % of Glc and 100 % of DGD for fraction F2 (containing only III; Table 2), respectively. This confirmed II to be a PI-DGD and III its Glc isomer PGlc-DGD. Compound IV was another major IPL of *T. barophilus* TLE (Figure 2, Table 1). Its molecular mass ([M+H]^+^ at *m/z* = 936.713), retention time (12.2 min), fragmentation pattern (for instance, ions at *m/z* = 138.1 and 204.1; Figure S2), and the presence of 52 % of Glc and 48 % of GlcNAc and of 100 % of DGD in fraction F1 (containing only IV) allowed to identify as a PGlcNAc-DGD (Table 2). The fragmentation pattern of IV did not allow to determine the exact position of the N-acetylation.

**Figure 2:**
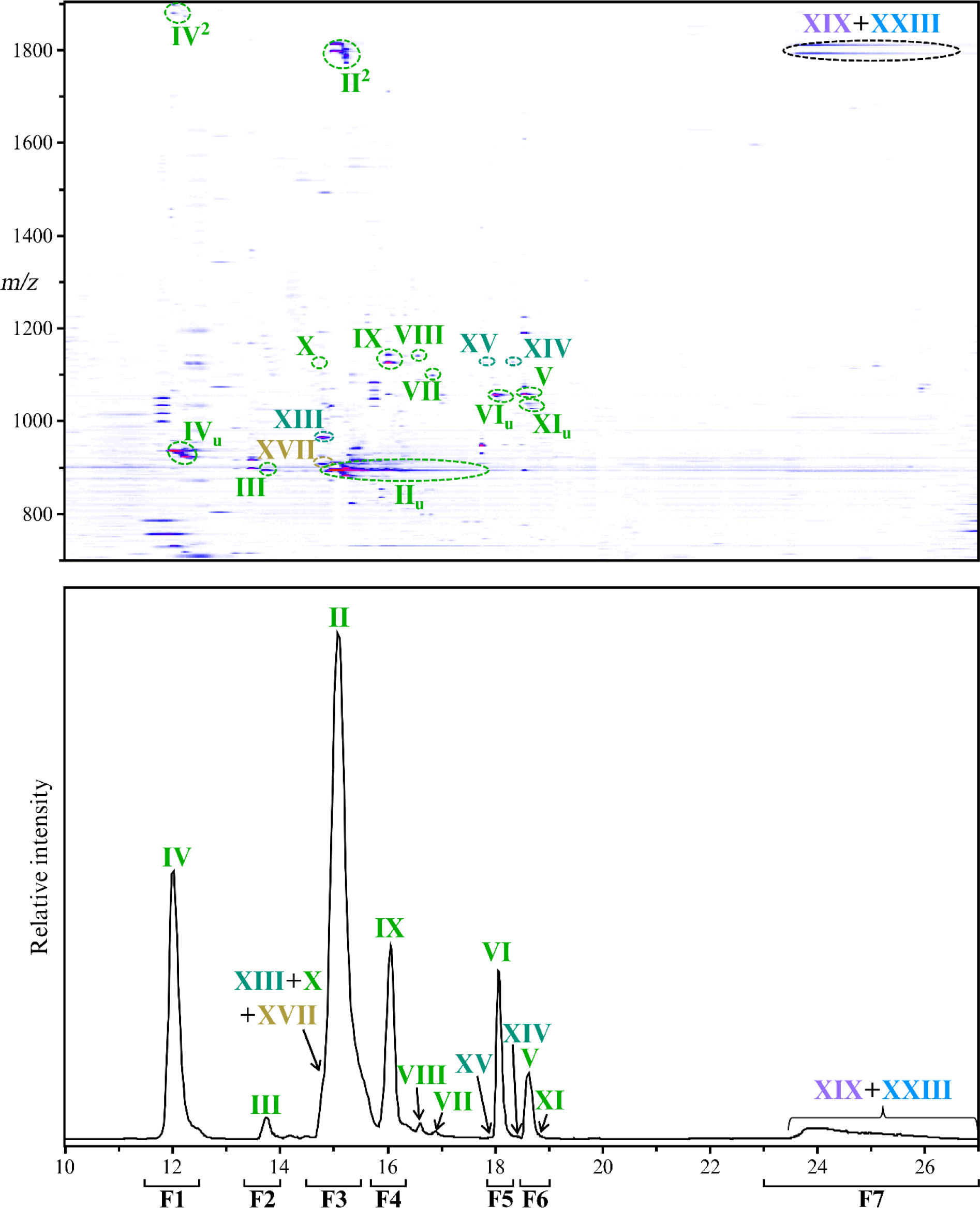
*Thermococcus barophilus* exhibits a large diversity of intact polar lipids. Intact polar lipids were detected in positive and negative ion mode. As no additional IPL could be identified in the negative ion mode, only the density map and chromatogram obtained in positive ion mode are displayed (zoom in the 10-27 min, *m/z* 700-1900 window). Compounds detected with unsaturations and/or diadducts are marked with _u_ and ^2^, respectively. The UHPLC chromatogram was drawn by extracting the following protonated ion masses with a mass deviation of ± 0.02 Da: 893.68, 894.70, 895.70, 909.72, 936.73, 965.78, 1038.69, 1056.77, 1057.75, 1098.78, 1124.83, 1125.82, 1126.85, 1127.83, 1139.81, 1786.36 and 1788.38. Refer to Figure 1, Table 1 and Figures S1-S9 for lipid structures and their respective molecular masses. F1-F7 delineate the time range corresponding to each fraction collected to confirm the structures of the identified lipids (refer to the Methods section).

**Figure 3:**
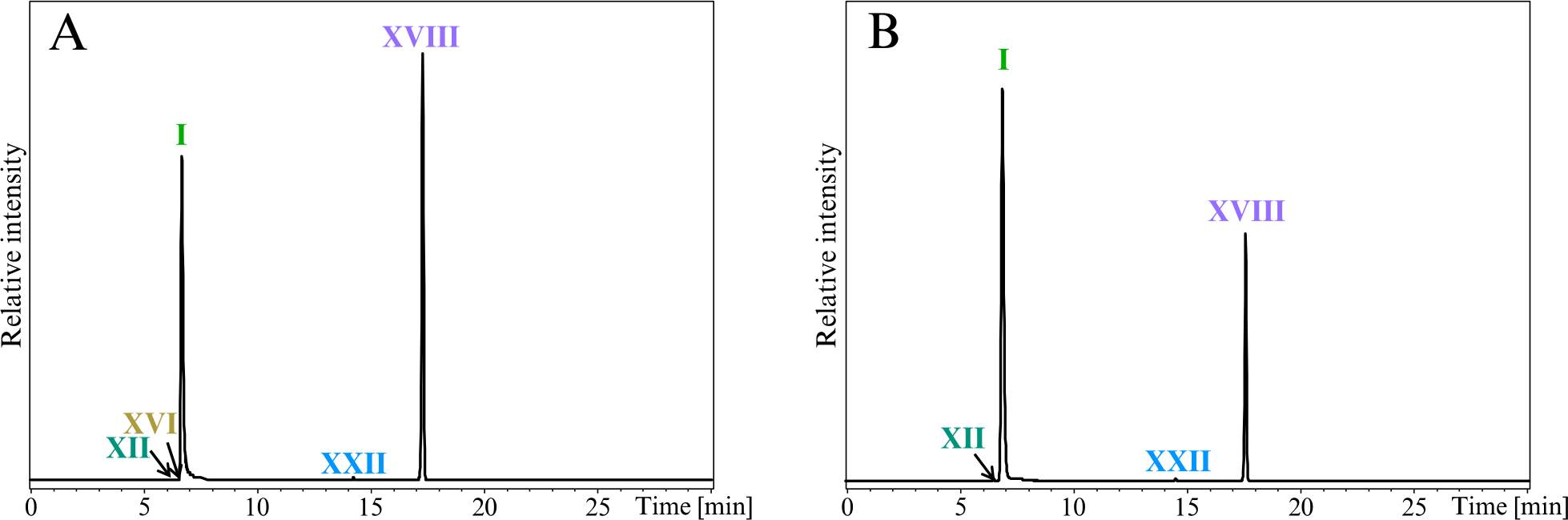
Analysis of the core lipids highlights a significant discrepancy between *Thermococcus barophilus* total and extracted lipids. Total CLs (**A**) and CLs from IPLs (**B**) were recovered after methanolysis of the biomass and the TLE, respectively. Direct methanolysis and intact polar lipid extraction were both performed on the same amount of biomass. UHPLC chromatograms were drawn in positive mode by extracting the following protonated ions with a mass deviation of ± 0.1 Da: 653.68, 667.70, 723.76, 743.71, 1302.32, 1304.34. Refer to Figure 1 for lipid structures.

**Table 1.**
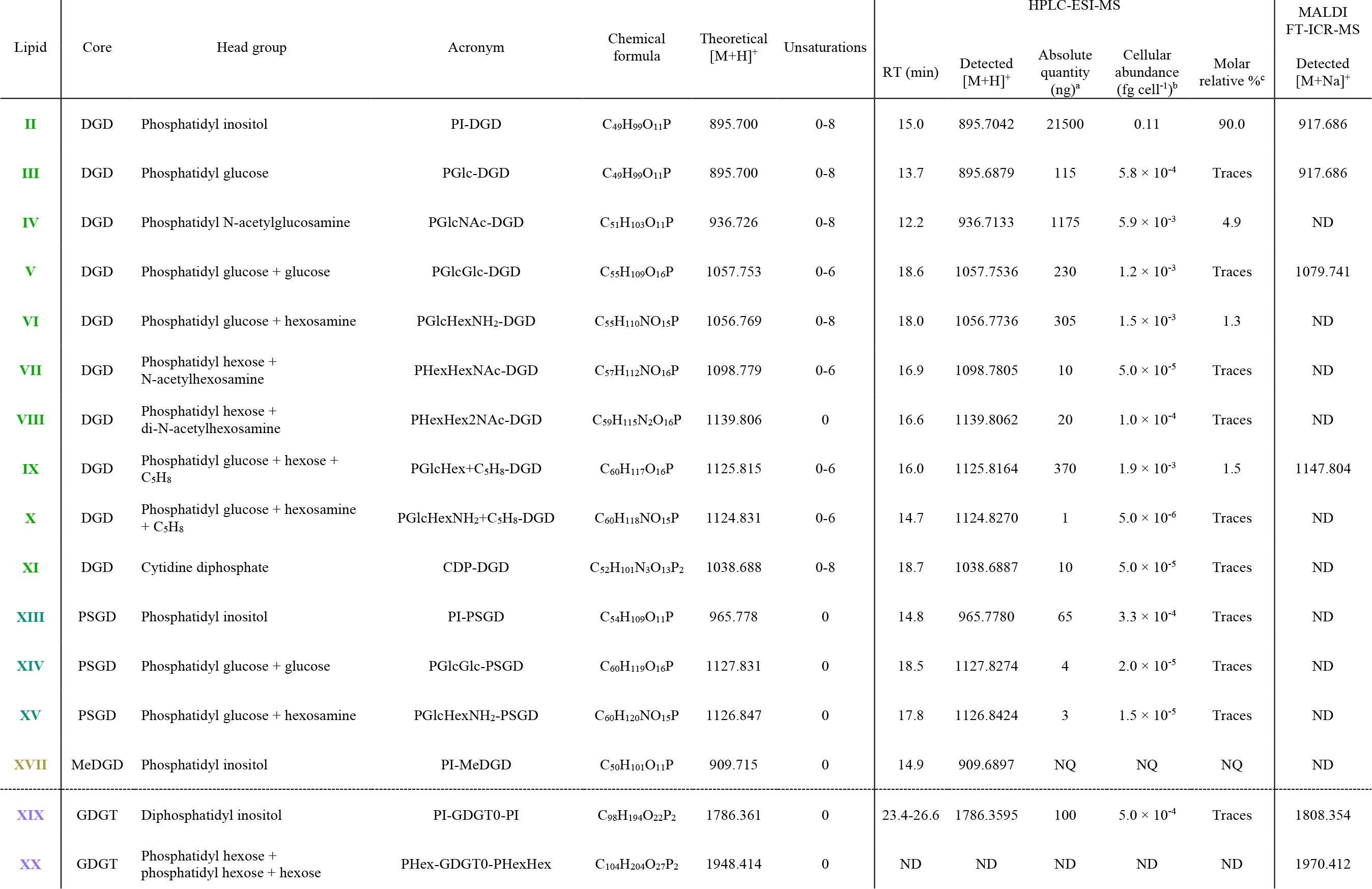
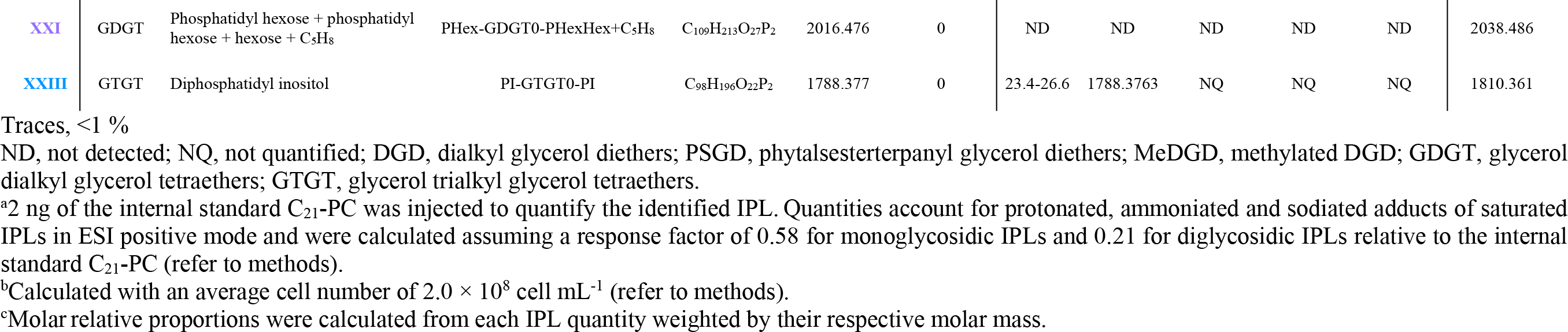
Intact polar lipid structures and lipid composition (absolute quantity, cellular abundance and molar relative %) of *Thermococcus barophilus*.

**Table 2.** Characteristics and distribution of the core structures and the polar head groups of purified major lipids of *Thermococcus barophilus*.

| Fraction | Time range (min) | Expected IPL | Detected IPL (%)<sup>a</sup> | Core structure<sup>b</sup> |  |  |  |  | Polar head<sup>c</sup> |  |  |  |  |  |  |  |  |
| --- | --- | --- | --- | --- | --- | --- | --- | --- | --- | --- | --- | --- | --- | --- | --- | --- | --- |
|  |  |  |  | DGD<br>**I** | PSGD<br>**XII** | MeDGD<br>**XVI** | GDGT0<br>**XVIII** | GTGT0<br>**XXII** | Pent | All/Fru | Man | Gal | Glc | Ino | 2Hex | GlcNAc | AcidoHex |
| F1 | 11.5-12.5 | **IV** | **IV** (100) | 100 | ND | ND | ND | ND | ND | ND | ND | ND | 58 | ND | ND | 42 | ND |
| F2 | 13.3-14.0 | **III** | **III** (100) | 100 | ND | ND | ND | ND | ND | ND | ND | ND | 100 | ND | ND | ND | ND |
| F3 | 14.5-15.5 | **II+X**<br>**+XIII+XVII** | **II**(98)+**XIII**(2)<br>**+XVII**(traces)+**X**(traces) | 98 | 2 | ND | ND | ND | ND | ND | ND | ND | ND | 100 | ND | ND | ND |
| F4 | 15.7-16.4 | **IX** | **IX**(93)+**II**(7)+**V**(traces)<br>**+XIII**(traces) | 100 | Traces | ND | ND | ND | ND | ND | ND | ND | 100 | ND | ND | ND | ND |
| F5 | 17.9-18.2 | **VI+XV** | **VI**(95)+**IX**(3)+**II**(2)<br>**+XV**(traces) | 100 | Traces | ND | ND | ND | ND | ND | ND | ND | 100 | ND | ND | ND | ND |
| F6 | 18.4-19.0 | **V+XI+XIV** | **V**(98)+**II**(1)+**VI**(1)<br>**+XIV**(traces) | 100 | Traces | ND | ND | ND | ND | ND | ND | ND | 100 | ND | ND | ND | ND |
| F7 | 23.0-27.0 | **XIX+XXIII** | **I** (100) | 2 | ND | ND | 95 | 3 | ND | ND | ND | ND | ND | 43 | 57 | ND | ND |

Traces, <1 %
ND, not detected; DGD, dialkyl glycerol diethers; PSGD, phytanylsesterterpanyl glycerol diethers; MeDGD, methylated DGD; GDGT0, glycerol dialkyl glycerol tetraethers with no cyclopentane ring; GTGT0, glycerol trialkyl glycerol tetraethers with no cyclopentane ring; Pent, β-D-xylofuranose, α-L-arabinopyranose and α-D-lyxopyranose; All/Fru, β-D-allopyranose and β-D-fructopyranose; Man, D-mannopyranose; Gal, β-D-galactopyranose; Glc, α-D-glucopyranose; Ino, myo-inositol; 2Hex, dihexoses; GlcNAc, $\alpha$ -N-acetyl-D-glucosamine; AcidoHex, D-galacturonic acid and D-glucuronic acid, D-galactosamine.
<sup>a</sup>Relative proportions account for protonated, ammoniated and sodiated adducts of saturated IPLs in ESI positive mode and were calculated assuming a response factor of 0.58 for monoglycosidic IPLs and 0.21 for diglycosidic IPLs relative to the internal standard C<sub>21</sub>-PC (refer to methods).
<sup>b</sup>Relative proportions account for protonated adducts in APCI positive mode and were calculated assuming a response factor of 0.42 for diethers and 0.57 for tetraethers relative to the internal standard C<sub>46</sub>-GTGT, or of 0.74 for diethers relative to tetraethers (refer to methods).
<sup>c</sup>Relative proportions were calculated assuming the response factors determined for each sugar using a standard solution (refer to methods).

Compounds XIII and XVII had retention times almost identical to compound II (14.8 and 14.9 min, respectively; Figure 2) but higher molecular masses ([M+H]^+^ at *m/z* = 965.778 and 909.690, respectively), and fragmentation patterns that suggested these compounds to be a phytanylsesterterpanyl glycerol diether (PSGD) and a methylated DGD (MeDGD) bearing phosphatidylhexose head groups (for instance, ions at *m/z* = 723.8 and 649.7, respectively; Figure S3). The analysis of fraction F3 containing compounds II, XIII and XVII showed exclusively Ino and 98 % of DGD and 2 % of PSGD (Table 2). The structures of PI-DGD II and PI-PSGD XIII were thus confirmed while compound XVII was only suggested to correspond to PI- MeDGD.

Compounds V and VI, which differed from one another by slightly less than one mass unit ([M+H]^+^ at *m/z* = 1057.754 and 1056.774, respectively) and showed similar fragmentation patterns (for instance, ions at *m/z* = 777.5 and 776.5, respectively; Figure S4), were identified as DGD bearing a glycosylated phosphatidylhexose head group and its hexosamine derivative in which one hydroxyl group is replaced by an amino group, respectively. The acid hydrolysis of the fractions containing V and VI (F6 and F5, respectively) yielded only Glc and DGD with traces of PSGD (Table 2). The presence of only Glc and the absence of disaccharides in both F5 and F6 suggested that the hydrolytic conditions were adequate to cleave off the glycosidic bond between the two sugar moieties, and that the structure of V was probably PGlcGlc-DGD. No hexosamine was detected in F5, impeding further characterization of compound VI, which we therefore tentatively assigned to PGlcHexNH_2_-DGD. Similarly to V and VI, compounds XIV and XV also differed from one another by slightly less than one mass unit ([M+H]^+^ at *m/z* = 1127.827 and 1126.842, respectively) and were present in fractions F6 and F5, respectively. Their molecular masses, shifted upwards by 70 mass units compared with PGlcGlc-DGD V and PGlcHexNH_2_-DGD VI, retention times (18.5 and 17.8 min, respectively), fragmentation patterns (for instance, ions at *m/z* = 685.5; Figure S5), and the presence of traces of PSGD in F6 and F5 (Table 2) suggested compounds XIV and XV to correspond to PGlcGlc-PSGD and PGlcHexNH_2_-PSGD, respectively. Similarly to compound VI, compounds VII, VIII, IX, and X ([M+H]^+^ at *m/z* = 1098.781, 1139.806, 1125.815, and 1124.831, respectively) could be associated to derivatives of PGlcGlc-DGD V (Table 1, Figures S4, S6, and S7). The comparison of the fragmentation patterns of VII and VIII with those of PGlcNAc-DGD IV and PGlcGlc-DGD V, and notably the detection of ions at *m/z* = 138.1 and 818.5 and at 245.1 and 859.5 (Figures S4 and S6), suggested the former to correspond to DGDs bearing mono- (PHexHexNAc-DGD VII) and di-N-acetylated glycosylated phosphatidylhexose (PHexHex2NAc-DGD VIII), respectively. The exact nature of the sugar moieties and the positions of the N-acetylations could not be further resolved. Compounds IX and X showed molecular masses ([M+H]^+^ at *m/z* = 1125.816 and 1124.827, respectively) shifted upwards by 68 mass units compared with PGlcGlc-DGD V and PGlcHexNH_2_-DGD VI, respectively, and displayed similar fragmentation patterns (for instance ions at *m/z* = 845.5 and 844.5; Figures S4 and S7). Although the nature of this increase in 68 Da could not be determined, we suggested it to be an isoprene unit (C_5_H_8_). The acid hydrolysis of fraction F4 (93 % of IX) released a majority of DGD and exclusively Glc (Table 2), and compound IX was thus assigned the partially solved structure PGlcHex+C_5_H_8_-DGD. Compound X was detected in very low amount (5.0 × 10^-6^ fg cell^-1^; Table 1), which hinders its complete characterization. Although no Glc was detected in F3 containing X, its similarity to both PGlcHexNH_2_-DGD VI and PGlcHex+C_5_H_8_-DGD IX (Figures S4 and S7) suggested it to correspond to PGlcHexNH_2_+C_5_H_8_. Other minor compounds with masses shifted upwards by either 68 or 70 Da were detected, e.g., a compound at 15.8 min with [M+H]^+^ at *m/z* = 1195.886 could correspond to PGlcGlc-PSGD with 68 additional Da, i.e., PGlcGlc+C_5_H_8_-PSGD. Their low abundances and the absence of MS^2^ spectra however prevented to solve their exact structures.

Apart from the ions typical of archaeal phospholipids (Figure S1), compound XI ([M+H]^+^ at *m/z* = 1038.689) showed a fragmentation pattern completely distinct from the IPLs described above (for instance, ions at *m/z* = 324.1 and 758.4; Figure S8) which suggested a head group distinct from typical hexoses. Compound XI could indeed be attributed to a DGD bearing a diphosphatidyl cytidine head group, i.e., the first intermediate in the pathway for polar head group fixation (47). The lack of relevant standard and the low quantities of XI prevented further elucidation of its structure by the acid hydrolysis of fraction F6 (Table 2).

In addition to their fully saturated forms, minute amounts of PI-DGD II, PGlc-DGD III, PGlcNAc-DGD IV, PGlcHexNH_2_-DGD VI, and CDP-DGD XI were detected with one to eight unsaturations whereas PGlcGlc-V, PHexHexNAc-VII, PGlcHex+C_5_H_8_- IX, and PHexHexNH_2_+C_5_H_8_-DGD X were detected with one to six unsaturations (Figure 2). Only the fully saturated forms were detected for the other diethers, namely PHexHex2NAc-DGD VIII, PI-PSGD XIII, PGlcGlc-PSGD XIV, PGlcHexNH_2-_PSGD XV and PI- MeDGD XVII.

As various Glc derivatives were detected, we looked for putative IPLs bearing similar and additional combinations of NH_2_, NAc and/or C_5_H_8_ groups, but none were detected. Similarly, glycolipids and phospholipids regularly found in Archaea, e.g., lipids bearing phosphatidylcholine (PC), phosphatidylserine (PS) and phosphatidylethanolamine (PE), were searched for but not detected in *T. barophilus*.

The very few tetraether-based IPLs detected in *T. barophilus* TLE all gather in a poorly resolved broad peak (Figure 2). The major component of this peak was compound XIX, whose molecular mass ([M+H]^+^ at *m/z* = 1786.360), fragmentation pattern (for instance, ions at *m/z* = 731.6 and 1544.3; Figure S9) and the acid hydrolysis of F7 (95 % of GDGT0, 43 % of Ino; Table 2) identified it as PI-GDGT0-PI. No fragmentation pattern was obtained for compound XXIII, but its molecular mass ([M+H]^+^ at *m/z* = 1788.376) shifted upwards by two mass units compared to XIX and the presence of 3 % of GTGT0 in F7 suggested that it might correspond to PI-GTGT0-PI (Table 2; see further details in Supplementary text). F7 also contained 57 % of a compound with the same fragmentation pattern as the Sac standard but with a slightly distinct retention time, which thus suggested it to be another disaccharide (hereafter referred to as 2Hex). The abundance of 2Hex in fraction F7 and the absence of tetraether-based IPLs with a 2Hex polar head group in UHPLC-ESI-MS suggested that this fraction might contain other, unresolved tetraether-based IPLs.

In addition to this set of IPLs, we detected a series of polyprenyl derivatives in *T. barophilus* TLE using our RP UHPLC method (Figure S11). These compounds represent a large family of membrane-bound polyisoprenoids known as major lipid carriers for membrane protein glycosylation in all three domains of life (48). Such compounds have been identified in a wide variety of archaea, in which the sugar residue can be attached to either alcohol, mono-, or diphosphate end-groups of polyprenyls with six to 14 isoprene units (49). Here, polyprenyl derivatives comprising 10 to 12 isoprene units and 1 to 5 unsaturations with only monophosphate head groups were identified (Figure S11).

### PI-DGD dominates the total lipid extract of *T. barophilus*

The IPLs identified in *T. barophilus* TLE were quantified considering a response factor of 0.58 for monoglycosidic (II, III, IV, XI, XIII, and XVII) and 0.21 for diglycosidic IPLs (V, VI, VII, VIII, IX, X, XIV, XV, XIX, and XXIII) relative to the internal standard C_21_-PC. Since unsaturated IPLs were detected in minute amounts, only the saturated forms were quantified. *T. barophilus* TLE (0.12 fg cell^-1^) was overwhelmingly dominated by PI-DGD II (91 %, 0.11 fg cell^-1^; Table 1), with a few other major IPLs, i.e., PGlcNAc-DGD IV, PGlcHexNH_2_-DGD VI, PGlcHex+C_5_H_8_-DGD IX, and PGlcGlc-DGD V (*ca.* 8 %, 1.05 × 10^-2^ fg cell^-1^). The remaining minor IPLs detected in trace amounts or in too low abundances to be quantified represented only 6.25 × 10^-3^ fg cell^-1^ (*ca.* 1 % of the TLE mass). DGD-based IPLs represented *ca.* 99 % of *T. barophilus* TLE, whereas tetraether-based IPLs were only recovered in trace amounts (Table 1).

To evaluate the efficiency of our extraction and analysis protocols, we performed several calculations to approximate a putative total IPL composition of TLE and of the original biomass. The acid methanolysis of the TLE provides reliable information on the IPL core lipid distribution (CLs from IPLs), which was composed of *ca*. 80 % of diethers and 20 % of tetraethers (Table 3). To appraise the lipid content per cell based on this CL composition, i.e., lipids without polar head group, an estimation of the nature and relative proportions of *T. barophilus* polar head groups is however required. The dedicated analysis of the polar head groups only allowed to access a few, low mass hexose derivatives (*ca.* 242 Da with a phosphatidyl group). Although none were detected here, a larger range of polar head groups, including smaller, e.g., PE (123 Da), and substantially larger ones, e.g., PHexHex2NAc (486 Da), sulfonotri- and tetrahexoses (551 and 713 Da), is to be expected in archaea (50, 51). This suggests that the polar head group can represent a non-negligible mass proportion of a given IPL, which we hypothesized to range from 0.2 (e.g., only small head groups like one and two PE for diether and tetraethers lipids, respectively) to 1 (e.g., only massive head groups like one and two phosphatidyltetrahexose for diether and tetraethers lipids, respectively) times the mass of the core lipid. Considering putative low (1:0.2) and high (1:1) mass ratios between the core lipid and the polar head group and the quantity of core lipids retrieved upon acid methanolysis (Table 3), we estimated that our TLE actually contained from 0.1 to 0.2 fg of lipids per cells. The 18 IPLs identified with UHPLC-MS (0.12 fg cell^-1^) thus represented 50-100 % of the TLE (Figure 4).

**Figure 4:**
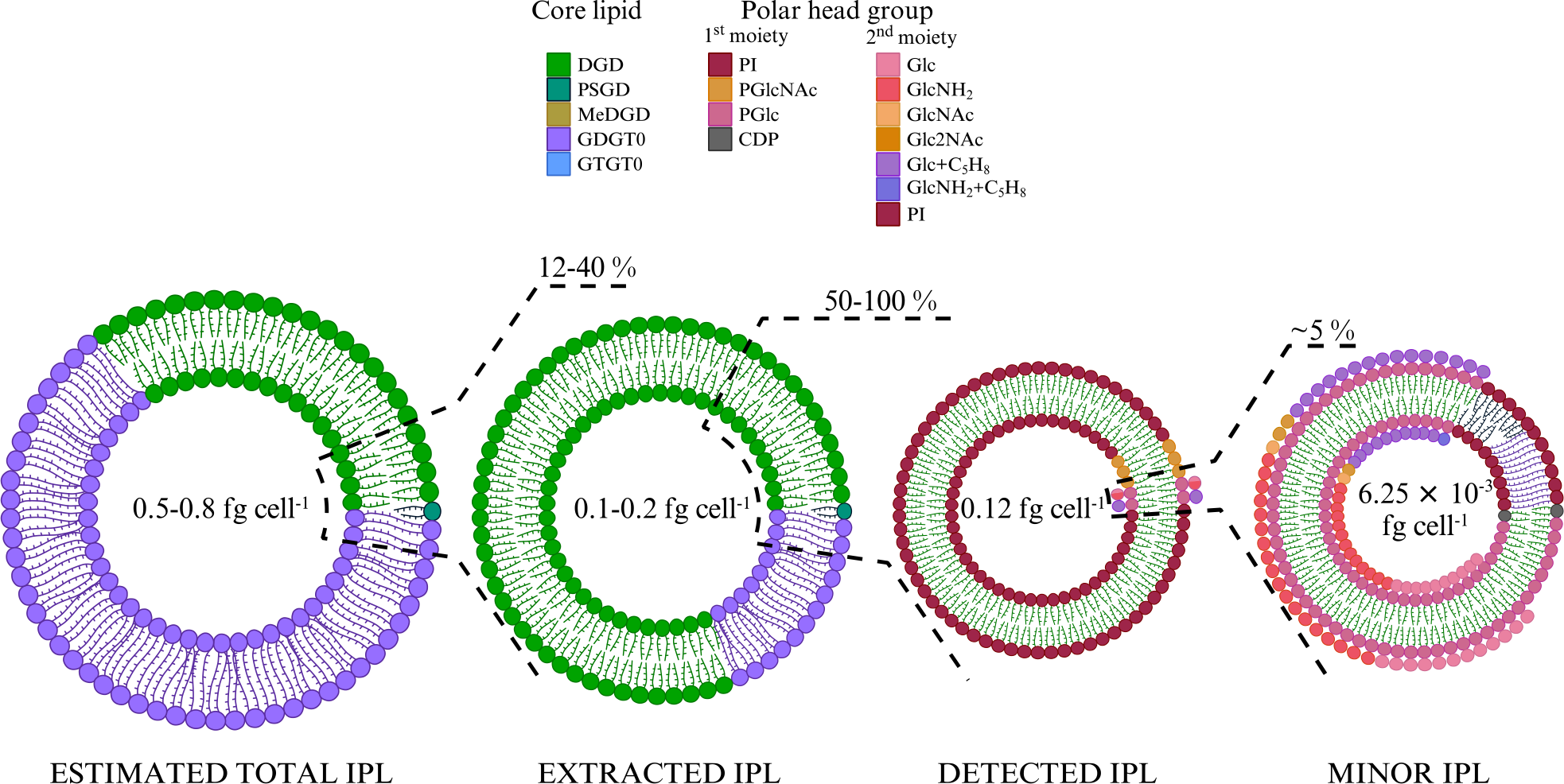
The vast majority of *Thermococcus barophilus*’ lipidome remains inaccessible. Pie chart representations of *Thermococcus barophilus* lipids at each extraction step. The direct acid methanolysis of *T. barophilus* biomass yielded its total core lipids content, which contained *ca*. 30 % of diethers (DGD, green; PSGD, dark green; MeDGD, dark yellow) and 70 % of tetraethers (GDGT0, purple; GTGT0, blue; Table 3). Considering putative low and high mass polar head groups found in Archaea, the direct acid methanolysis of the biomass yielded an experimentally-derived theoretical lipid content of 0.5 to 0.8 fg of lipids cell^-1^. Acid methanolysis of the TLE allowed the determination of the extraction yield on *T. barophilus* (0.1 to 0.2 fg cell^-1^ considering putative low and high mass polar head groups, *ca.* 12-40 % of the biomass’ total lipids) and of the diether and tetraether distribution in extracted lipids (80/20; Table 3). Although part of *T. barophilus* IPLs is indeed extracted, the majority of its IPLs, and especially tetraether- based ones, remain resistant to extraction. 18 IPLs representing 50-100 % of these extracted lipids (0.12 fg cell^-1^) were identified with UHPLC-MS (Table 1). The putative discrepancy between extracted and identified IPLs highlights only a partial detection of *T. barophilus* IPLs with our UHPLC-MS system. Complex IPLs with at least two sugar residues (1^st^ and 2^nd^ polar head moiety) were recovered with an even lower yield, i.e., 6.25 × 10^-3^ fg cell^-1^, *ca.* 5 % of the IPLs identified (Table 1), thus showing that peculiar polar head groups might be the main reason for the detection defect observed in *T. barophilus*.

**Table 3.** Core lipid composition of the biomass (totCLs) and of the total lipid extract (CLs from IPLs) of *Thermococcus barophilus*.

|  | Diethers<sup>a</sup> |  |  |  |  |  | Tetraethers<sup>a</sup> |  |  |  |  |  | Total |  |
| --- | --- | --- | --- | --- | --- | --- | --- | --- | --- | --- | --- | --- | --- | --- |
|  | DGD **I** |  |  | PSGD **XII** |  |  | GDGT0 **XVIII** |  |  | GTGT0 **XXII** |  |  |  |  |
|  | μg | fg cell<sup>-1</sup> | Rel% | μg | fg cell<sup>-1</sup> | Rel% | μg | fg cell<sup>-1</sup> | Rel% | μg | fg cell<sup>-1</sup> | Rel% | fg cell<sup>-1</sup> | D/T |
| TotCLs | 22.2 | 0.1 | 28 | 6.1 × 10<sup>-2</sup> | 3.1 × 10<sup>-4</sup> | Traces | 56.8 | 0.3 | 72 | 0.4 | 2.2 × 10<sup>-3</sup> | Traces | 0.4 | 0.39 |
| CLs from IPLs | 14.6 | 7.3 × 10<sup>-2</sup> | 80 | 4.2 × 10<sup>-2</sup> | 2.1 × 10<sup>-3</sup> | Traces | 3.6 | 1.8 × 10<sup>-2</sup> | 20 | 3.1 × 10<sup>-2</sup> | 1.5 × 10<sup>-4</sup> | Traces | 9.1 × 10<sup>-2</sup> | 4.0 |

DGD, dialkyl glycerol diethers; PSGD, phytanylsesterterpanyl glycerol diethers; GDGT0, glycerol dialkyl glycerol tetraethers with no cyclopentane ring; GTGT0, glycerol trialkyl glycerol tetraethers with no cyclopentane ring; D/T, diethers over tetraethers ratio; Rel%, molar relative proportion; μg, absolute lipid quantity in μg; fg cell<sup>-1</sup>, cellular lipid abundance in fg cell<sup>-1</sup> calculated with an average cell number of 2.0 × 10<sup>8</sup> cell mL<sup>-1</sup>; ND, not detected. Traces, <1 % <sup>a</sup>Relative proportions account for protonated adducts in APCI positive mode and were calculated assuming a response factor of 0.42 for diethers and 0.57 for tetraethers relative to the internal standard C<sub>46</sub>-GTGT, or of 0.74 for diethers relative to tetraethers (refer to methods).

To further estimate how close this lipid composition was from the real lipidome of *T. barophilus*, we compared it with calculated and an experimentally-derived theoretical total lipid contents (refer to Figure 4 for a summary of these estimations). *T. barophilus* cells are coccoid with a cell diameter of 0.8 to 2.0 µm (36) and covered in one or more dense proteinaceous surface layers like other Thermococcales (52). Using the calculations described by Lipp *et al.* (31) with a membrane thickness of 5.5 nm and protein content of 70 %, our calculated theoretical total lipid content ranged between 3.0 and 20 fg cell^-1^. On the other hand, the direct acid methanolysis of *T. barophilus* biomass yielded its total core lipid content (total CLs), which contained *ca*. 30 % of diethers and 70 % of tetraethers (Table 3). As for the estimation of the TLE content, considering low and high mass ratios between the core lipid and the polar head group resulted in an experimentally-derived theoretical total lipid content of *T. barophilus* ranging from 0.5 to 0.8 fg cell^-1^. Our TLE (0.1 to 0.2 fg cell^-1^) thus represented *ca*. 12-40 % of *T. barophilus* lipidome (experimental-derived theoretical lipid content; Figure 4).

### MALDI-FT-ICR-MS reveals novel tetraether-based IPLs in *T. barophilus*

MALDI-FT-ICR-MS allows for the direct ionization of a sample, without prior wet-chemical treatment, and thus offers a valuable tool to explore archaeal IPLs that are unreachable by extraction-based analytical procedures. The direct ionization of *T. barophilus* biomass using MALDI-FT-ICR-MS resulted in clusters of peaks for each IPL (Figure 5), suggesting the presence of numerous isotopologues and singly charged adducts, i.e., [M+H]^+^, [M+Na]^+^, [M+K]^+^, [M+2Na-H]^+^, [M+K+Na-H]^+^ and [M+3Na-2H]^+^. Lipids were assigned based on the molecular masses of the different adducts detected, the monosodiated one being always the most abundant. Under the optimal experimental conditions determined here (not shown), we could detect the majority of *T. barophilus* most abundant diether-based IPLs identified in UHPLC-MS, i.e., PI-DGD II and/or PGlc-DGD III, PGlcGlc-DGD V and PGlcHex+C_5_H_8_-DGD IX, although some were not observed, e.g., PGlcNAc-DGD VI. A special focus was paid to specifically target *T. barophilus* unidentified tetraether lipid diversity, and PI-GDGT0-PI XIX and PI-GTGT0-PI XXIII indeed appeared in a seemingly much higher abundance than that revealed by UHPLC-ESI-MS (Figure 5). Other, lower mass ion adducts were detected, e.g., at *m/z* = 1646.304, 1566.339 and 1324.390, and were assigned to sodiated adducts of PI-GDGT0-P, PI-GDGT0 and GDGT0 XVIII, respectively. As those compounds were not identified in *T. barophilus* TLE analyzed with UHPLC-ESI-MS, they were assumed to result from laser-induced partial degradation of the much more abundant PI-GDGT0-PI XIX (Figure 5). Additionally, two putative uncharacterized tetraether-based IPLs were observed, namely compounds XX and XXI. Compound XX showed molecular ions at *m/z* = 1948.451, 1970.413 and 1992.396 that could be assigned to the [M+H]^+^, [M+Na]^+^ and [M+2Na-H]^+^ adducts of PHex-GDGT0-PHexHex (theoretical ions at *m/z* = 1948.414, 1970.396 and 1992.378, respectively). Similarly, molecular ions of compound XXI at *m/z* = 2016.483, 2038.485 and 2060.468 were assigned to [M+H]^+^, [M+Na]^+^ and [M+2Na-H]^+^ adducts of PHex-GDGT0- PHexHex+C_5_H_8_ (theoretical ions at *m/z* = 2016.476, 2038.458 and 2060.440, respectively). Despite a relatively large gap between theoretical and observed masses which might result from very low amounts of those compounds and the increasing absolute mass error with higher masses, these putative structures were further supported by the detection of 2Hex polar head groups in fraction F7 corresponding to the tetraether unresolved peak in UHPLC-MS (Table 2). Other tetraether-based IPLs bearing polar head groups similar to those detected on the diether-based IPLs, such as (poly)N-acetylated hexosamine, were also screened for but not detected. Scanning *T. barophilus* biomass for higher masses neither yielded other ions nor improved the recovery of the newly identified IPLs (not shown).

**Figure 5:**
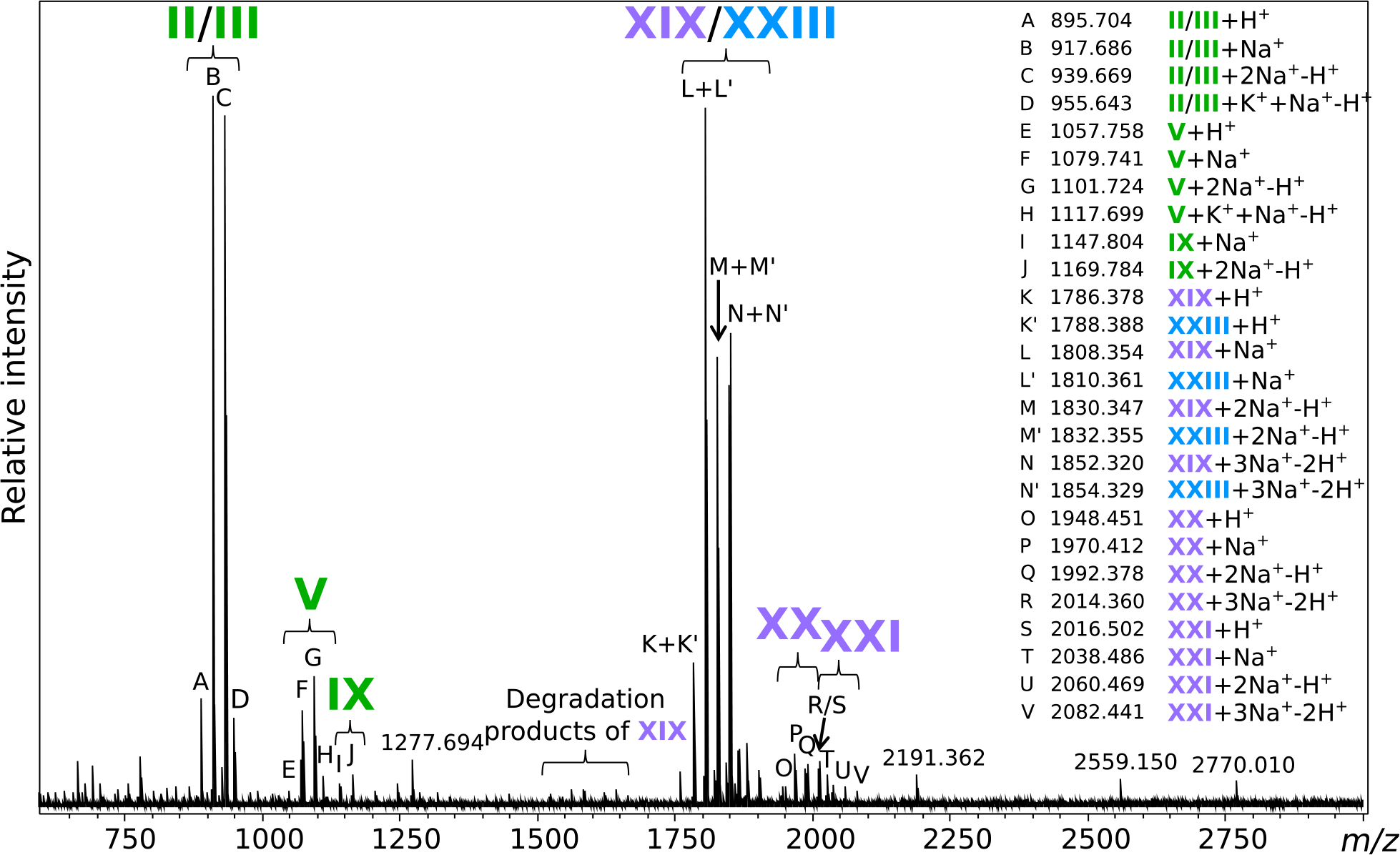
Extraction-free analysis reveals novel tetraether-based intact polar lipids in *Thermococcus barophilus*. *m/z* detected for each marked peak and the putative corresponding lipid adduct are listed on the right side of the figure. Masses similar to that expected from partial hydrolysis of tetraethers, e.g., PI-GDGT0-P, PI- GDGT0 and P-GDGT0, are indicated as degradation products of PI-GDGT0-PI XIX. The masses of major unidentified peaks are also displayed. Refer to Figure 1 for lipid structures.

Altogether, 18 saturated and 64 unsaturated IPLs were detected and tentatively identified in *T. barophilus*. Structures of PI-DGD II, PGlc-DGD III, PGlcNAc-DGD IV, PGlcGlc-DGD V, PGlcHexNH_2_- DGD VI, PGlcHex+C_5_H_8_-DGD IX, PI-PSGD XIII, PGlcGlc-PSGD XIV, PGlcHexNH_2_-PSGD XV, and PI-GDGT0-PI XIX were validated by analyzing independently their polar head groups and core structures, whereas those of PHexHexNAc-DGD VII, PHexHex2NAc-DGD VIII, CDP-DGD XI, and PI-MeDGD XVII were determined based on their fragmentation patterns alone. In contrast, the structure of PI-GTGT0- PI XXIII derived solely from the acid hydrolysis results, whereas those of PHex-GDGT0-PHexHex XX and of PHex-GDGT0-PHexHex+C_5_H_8_ XXI resulted from the molecular masses detected by FT-ICR-MS.

## Discussion

### Novel IPL structures were uncovered from the diverse lipid composition of *T. barophilus*

By means of UHPLC-MS and MALDI-FT-ICR-MS, 18 saturated and 64 unsaturated IPLs were identified in *T. barophilus* (Table 1). Fourteen IPLs were based on diethers, i.e., DGD I, PSGD XII and MeDGD XVI, and four on tetraethers, i.e., GDGT0 XVIII and GTGT0 XXII. Ten distinct polar head groups were detected, among which three were derivatives of phosphatidylhexose (PI, PGlc, PGlcNAc), six of glycosylated phosphatidylhexose (PGlcGlc, PGlcHexNH_2_, PHexHexNAc, PHexHex2NAc, PGlcHex+C_5_H_8_, PHexHexNH_2_+C_5_H_8_), and one of nucleoside diphosphate (CDP; Table 1).

Similarly to numerous other archaea, and especially Thermococcales (35, 42, 53–55), PI-DGD II was the dominant IPL of *T. barophilus*. Whereas PSGD-based IPLs were reported in numerous halophilic archaea and a few methanogens (see for instance (22, 56–58)), this study reported for the first time the presence of PSGD and MeDGD-based IPLs in Thermococcales and in hyperthermophilic archaea. In addition to *T. barophilus*, PI-GTGT0-PI XXIII has been reported in *Thermococcus kodakarensis*, *Pyrococcus furiosus* and *P. yayanosii* (40) and may be a common IPL to all Thermococcales, as the core lipid GTGT0 XXII was reported in every Thermococcales investigated so far (37). Glc has been reported repeatedly as a major sugar residue in archaeal glycolipids (see, for example, (59–63)), but only rarely in phosphoglycolipids, i.e., in *Aeropyrum pernix* (64) and in *Thermococcus zilligi* (54), a close relative of *T. barophilus*. Meador *et al*. (35) recently updated the IPL composition of *Thermococcus kodakarensis*, and notably identified PHexNAc-DGD, PHexHex-DGD, PHexHexNH_2_-DGD and PHexHexNH_2_+C_5_H_8_-DGD, with no further characterization of the polar head groups. To our knowledge, our detailed investigation of the sugar residues is the first report of such a diversity of Glc derivatives as polar head groups of phosphoglycolipids in Archaea, which were initially assumed to be mostly built upon Ino. This study also reported for the first time mono- and diacetylated PHexHex-DGD VII and VIII as well as PHexHex+C_5_H_8_ as a GDGT0 polar head group in XXI. Although our results do not drastically contrast with the lipid composition of other Thermococcales, they extend the known lipid diversity for this order of Archaea and beyond, and places *T. barophilus* as a prime model for further investigation of the Thermococcales membrane composition, organization and adaptation.

### A combination of UHPLC-MS and MALDI-FT-ICR-MS to elucidate *T. barophilus* IPL composition

Archaeal lipid extraction and fractionation have previously been demonstrated to be biased towards certain lipid classes (24, 30, 65). The first and only description of *T. barophilus* intact polar lipids, which reported exclusively PI-DGD II (36), undoubtedly suffered from such biases. The reevaluation of *T. barophilus* CLs indeed showed an abundance of tetraether-based IPLs and demonstrated the impossibility to exhaustively extract its IPLs with typical extraction procedures (24). While the reassessment of *T. barophilus* IPLs in the present study did provide a greater insight into its diversity, including tetraether-based IPLs (Table 1), we could only access 0.12 fg of lipid cell^-1^. Although this remains in line with observations made for other archaea, e.g., *T. barophilus* close relative *T. kodakarensis* (0.38 to 1.61 fg cell^- 1^ (35)), our results highlighted three limitations of the current procedure that can explain such a low yield.

First, we observed a large difference between *T. barophilus* calculated and experimentally-derived theoretical lipid contents (3.0 to 20.0 vs. 0.5 to 0.8 fg cell^-1^), which suggests that our calculations might be far off *T. barophilus* real total lipid content. Indeed, even adding up the other lipid molecules *T. barophilus* synthesizes and that are not visible after methanolysis of the biomass, i.e., polyprenyl phosphates (Figure S10) and apolar polyisoprenoids (up to 1 % of the membrane content, (24)), to our experimentally-derived theoretical total lipid content would not be enough to reach the calculated theoretical total lipid content. The model for cell lipid content, initially built for bacterial cells and used here with rough estimates of *T. barophilus* cell diameter and membrane lipid/protein ratio, thus requires revisions to better fit the archaeal cell membrane. Although *T. barophilus* experimentally-derived theoretical total lipid content might also be inaccurate, for instance due to other, unidentified polar head groups larger than the phosphatidylmono- and di-hexoses considered here, it was used hereafter as a reference (Figure 4). Although closer to the total IPLs detected, the experimentally-derived theoretical total lipid content still showed a large discrepancy with what we could actually access with our B&D extraction (0.5 to 0.8 vs. 0.12 fg cell^-1^), suggesting that other biases might hinder the elucidation of *T. barophilus* entire lipidome.

We observed a major discrepancy between this experimentally-derived theoretical value and extracted lipid contents (0.5 to 0.8 vs. 0.1 to 0.2 fg cell^-1^; Figure 4), which indicates that most of *T. barophilus* lipids are resistant to our extraction procedure. Our TLE contained only *ca*. 20 % of tetraethers (Table 3), whereas they represented from 45 to 70 % of *T. barophilus* experimentally-derived theoretical total lipid content in previous studies (24, 37) and here, respectively. Despite inconsistencies in the total amount of tetraethers *T. barophilus* synthesizes that might stem from different growth or extraction/analytical conditions, all three studies agree on a large discrepancy of diether/tetraether distributions between the experimentally-derived theoretical total lipid content and the TLE. This suggests that most of the IPLs not recovered are built upon tetraethers, and thus supports a major deficiency of our extraction procedure in recovering tetraether-based IPLs.

We also highlighted a minor inconsistency between extracted and detected IPLs (0.1 to 0.2 vs. 0.12 fg cell^-1^), which suggests that part of the IPLs that are indeed extracted might remain invisible to our detection method. Despite being composed of up to 20 % of tetraethers (Table 3), *T. barophilus* TLE displayed only two tetraether-based IPLs upon UHPLC-ESI-MS analysis, i.e., PI-GDGT0-PI XIX and PI- GTGT0-PI XXIII, which represented less than 1 % of the TLE (Table 2). Although a fraction of tetraether-based IPLs are indeed extracted and present in the TLE, these results suggest that they remain resistant to detection by our analytical setup. Additionally, *T. barophilus* TLE was overwhelmingly dominated by PI- bearing IPLs both in this study (90 %, Table 1, Figure 4) and that of Marteinsson *et al.* (100 %) (36), while neither glycolipids nor other polar head groups typically found in Archaea (e.g., PE, PS and PG, (15, 35, 50, 60, 66)) were observed. It is now widely accepted that physicochemical properties and physiological and adaptive functions of biological membranes are governed by the structural diversity of both the alkyl chains and the polar head groups found in the lipidome (67). One may thus speculate that a natural membrane containing almost exclusively one polar head group might not be biologically functional. While the absence of typical archaeal IPLs in *T. barophilus* might be linked to its particular membrane physiology and/or environmental conditions, a functional membrane composed of > 90 % of a single IPL is hardly conceivable, and *T. barophilus* membrane should theoretically contain other isomers and derivatives of phosphatidyl(poly)hexoses not detected here to be functional. This therefore suggests that our UHPLC-ESI-MS analytical procedure, and probably our B&D extraction method as well, might artificially enhance the detection of PI-based over other phosphatidylhexose-derivative IPL populations. Altogether, our results thus highlight two major shortcomings of our extraction and analytical procedure, i.e., preferential extraction and detection of 1) diether-based and 2) PI-bearing IPLs, which resulted in a *T. barophilus* lipidome artificially composed of almost exclusively PI-DGD II. However, our study shows that there is still much to explore in the lipidome of *T. barophilus* and provides clues about the presence of other lipids such as tetraethers with derivatives of phosphatidyl(poly)hexoses probably based on distinct sugar moieties.

In contrast to UHPLC-MS, MALDI-FT-ICR-MS of *T. barophilus* cell pellet revealed high levels of several tetraether lipids when focusing on high *m*/*z*. For instance, PI-GDGT0-PI XIX appeared as one of *T. barophilus* main IPLs with our MALDI-FT-ICR-MS settings (Figure 5). In addition to PI-GDGT0-PI XIX, other low-mass tetraether derivatives such as PI-GDGT0-P, PI-GDGT0 and P-GDGT0 were identified (Figure 5). PI-GDGT0 has been repeatedly reported in Archaea, including Thermococcales (35, 40, 68), but almost always using rather destructive ionization methods (for instance, fast-atom bombardment in (68)). Similarly, PI-GDGT0 was only detected here with MALDI-FT-ICR-MS under the highest laser power setting (not shown) and not with our soft UHPLC-ESI-MS method (Figure 2). This suggests that our MALDI-FT-ICR-MS procedure might alter tetraether lipids, preventing their detection as IPLs, and that the aforementioned compounds could stem from laser-induced degradation rather than be true IPLs of *T. barophilus*. MALDI-FT-ICR-MS nonetheless allowed to access previously unknown tetraether-based IPLs, i.e., PHex-GDGT0-PHexHex XX and PHex-GDGT0-PHexHex+C_5_H_8_ XXI. Altogether, these results proved MALDI-FT-ICR-MS to be a prime alternative to UHPLC-MS for exploring archaeal lipid diversity, and especially tetraether-based IPLs. In the future of MALDI-FT-ICR-MS lipidomics, fine-tuning of the laser parameters, including laser-based post-ionization (69), and of the matrix composition should help access an even wider archaeal IPL diversity, although combination with UHPLC-MS remains necessary for lipid quantitation and to elucidate the complete lipidome of Archaea.

### Insights into *T. barophilus* membrane organization

Most of the physical properties of archaeal lipids were based on the study of synthetic PE- and PC-bearing lipids, and the absence of a comprehensive inventory of *T. barophilus*’ polar head groups prevented further understanding of its membrane physicochemical properties and organization. The detection of 82 IPLs however opens new avenues for understanding the membrane contribution and biological functions of archaeal polar head groups as even the least abundant lipids were shown to ensure key cellular functions in both Bacteria and Archaea (70, 71).

PI, present in PI-DGD II, PI-PSGD XIII, PI-MeDGD XVII, PI-GDGT0-PI XIX and PI-GTGT0-PI XXII, was the most abundant polar head group in *T. barophilus* TLE (Table 1). Physical studies using neutron and x-ray diffractions and NMR spectrometry demonstrated that PI extended deeply into the aqueous environment in a slightly tilted configuration relative to the membrane surface normal (72–74). The extension of PI away from the membrane surface favors intra- and intermolecular hydrogen bonds, thus creating an extensive network that shields the membrane with a large electrostatic barrier preventing proton and ion leakages (26, 72, 74). In addition, the direct projection of PI into the aqueous environment allows for a maximum hydration of the inositol ring (74), which turns into a bulky hydrated head group. The high volume of this bulky head group relative to that of the lipid alkyl chains enhances the conformational freedom of the latter, which might eventually result into a looser packing and a higher water permeation in model membranes containing PI (75). This packing defect generated by the PI head group, especially in the tightly packed archaeal isoprenoid membranes, would in turn unlock more loading space for membrane proteins and the higher water permeation would allow for solvent interactions essential to protein stability within the membrane environment (75, 76). In contrast, the bulky head group creates a repulsive hydrated layer that stabilizes the membrane by preventing deformation and membrane fusion (77, 78). The various membrane macrostructures observed in Thermococcales, e.g., nanotubes and vesicles (79, 80), should theoretically be greatly disfavored if their membrane was indeed composed exclusively of PI-based lipids that prevent membrane remodeling. These results further confirm our assumption that PI might not necessarily be the major head group in *T. barophilus* despite its overwhelming dominance in our extracts (Table 1). In contrast, low proportions of PI would provide an enhanced fluidity in an otherwise tightly packed archaeal membrane while preserving its impermeability.

In addition to inositol, glucose was found in a variety of polar head group derivatives that represented *ca.* 8 % of *T. barophilus* TLE (Table 1). Due to the structural similarities between hexose isomers, one might speculate that their effects on biological membranes would be comparable but synthetic glycolipids bearing distinct hexose moieties showed different orientations relative to the membrane surface. Glc nonetheless displayed an extended conformation similar to that of Ino (81), suggesting that it might support membrane physicochemical properties analogous to those described above.

Stereochemical changes of a single hydroxyl group were shown to dramatically alter membrane properties and stability (9, 82). Various derivatives of Glc bearing distinct additional groups were identified in *T. barophilus* (Table 1), but their exact position on the hexose ring and the presence of different position isomers could not be ascertained although they might support distinct functions. For instance, positions of the phosphatidyl groups in phosphoinositides impacted their ionization properties, hydrogen bond networks and thus their interactions with membrane proteins and lipids (83). No data are currently available on the alterations of the lipid properties generated by the additional NAc, NH_2_ and C_5_H_8_ moieties detected in *T. barophilus*. However, an O-acetylation on the C-6 atom of the Glc ring of a fatty-acyl analogue of PGlc- DGD III found in various bacterial and mammalian cells was demonstrated to change the immunogenic properties of the IPL (84, 85), suggesting that NAc-bearing IPLs of *T. barophilus* might exert different properties than their hydroxylated forms. Based on the polarity of these moieties, one might also speculate that NAc and NH_2_ would behave similarly than the regular hydroxyl group, whereas the apolar C_5_H_8_ could cause dramatic changes in the orientation, hydration and interaction of the monosaccharide ring.

The polar moiety of lipids bearing diglycosides have been shown to extend away from the surface and to generate intermolecular hydrogen bonds, hence conferring the diglycosidic lipids similar physicochemical properties than those of monoglycosidic ones (86–88). In addition, the even higher relative volume of the polar head group might result in further enhanced conformational freedom of the alkyl chain. Derivatives of glycosylated phosphatidylhexose lipids, such as PHexHex, PHexHexNH_2_ and PHexHex+C_5_H_8,_ have been detected in Bacteria, Eukarya and Archaea (35, 89). Although their biological purpose remains elusive in Archaea, these lipid derivatives notably act as protein anchor in bacterial and eukaryotic membranes (90), and similar functions might be expected for their archaeal counterparts.

Altogether, these results enhance our comprehension of the membrane structuration suggested for *T. barophilus* and highlight putative biological functions for the different IPLs detected in this study. Characterization of the physicochemical properties of synthetic or natural archaeal lipids with phosphoglycosidic head groups remains nonetheless essential to precisely define the role of the diverse archaeal lipid compositions in membrane physiology and organization.

## Conclusions

We reassessed here the intact polar lipid, core lipid and lipid polar head group compositions of *Thermococcus barophilus*, a model for membrane architecture and adaptation to extreme conditions in Archaea. We unraveled the presence of at least 82 distinct membrane lipids, including a variety of core structures, i.e., saturated and unsaturated DGD, MeDGD, PSGD, GDGT and GTGT, and the widest diversity of polar head groups in Thermococcales known to date. Although not drastically different from that of *T. barophilus* close relatives, the lipid composition reported here extends the known diversity of phosphoglycosidic head groups known in Thermococcales. In agreement with previous investigations of *T. barophilus* and other Thermococcales IPLs, the lipid diversity revealed here was overwhelmingly dominated by PI-DGD. The low extraction yield, the excessive prevalence of PI-DGD, the low diversity of polar head group moieties compared to bacterial and eukaryotic lipidomes (although high compared to other archaeal lipidomes) and the CL released upon acid methanolysis demonstrated that a large portion of *T. barophilus* lipidome still remains inaccessible to the employed extraction and analytical protocols. Extraction-free analysis with MALDI-FT-ICR-MS allowed to access previously undetected tetraether-based IPLs, and further improvements and developments of new methodologies might pave the way to the discovery of completely new archaeal IPLs. Due to the isoprenoid alkyl chains of archaeal lipids, *T. barophilus* membrane is tightly packed, but the addition of bulky phosphatidylhexose head groups might provide relaxation while maintaining impermeability. Altogether, our results illustrate the complexity and diversity of *T. barophilus* membrane structure and pose this species as a prime model to elucidate archaeal membrane lipid diversity, properties and organization.

## Acknowledgments

M.T. is supported by a Ph.D. grant from the French Ministry of Research and Technology. The authors would like to thank the French National Research Agency for funding the ArchaeoMembranes project (ANR-17-CE11-0012-01) and the CNRS Interdisciplinary program ’Origines’ for funding the ReseArch project. Research at MARUM was funded by Germany’s Excellence Strategy (EXC-2077) project 390741603 “The Ocean Floor – Earth’s Uncharted Interface”.

## Author contributions

Conceptualization, funding acquisition, project administration and supervision, K-U.H. and P.M.O.; Formal analysis, M.T. and S.C.; Investigation, M.T., S.C., and L.W.; Methodology, M.T., S.C., J.S.L. and L.W.; Visualization and writing – original draft, M.T.; Writing – Review and editing, M.T., S.C., L.W., J.S.L, K-U.H. and P.M.O.

## Abbreviations

IPL: intact polar lipid(s)
CL: core lipid(s)
MGD: monoalkyl glycerol diethers
DGD: dialkyl glycerol diethers
PSGD: phytanylsesterterpanyl glycerol diethers
MeDGD: methylated DGD
GDGT: glycerol dialkyl glycerol tetraethers
GTGT: glycerol trialkyl glycerol tetraethers
MGDG: monogalactosyl diacylglycerol
DGDG: digalactosyl diacylglycerol
C_46_-GTGT: GTGT containing 46 carbon atoms
PI: phosphatidyl inositol
PHex: phosphatidyl hexose
PHexNAc: phosphatidyl N-acetylhexosamine
PHexHex g: lycosylated phosphatidyl hexose
PHexHexNH_2_: hexosamine phosphatidyl hexose
PHexHexNAc: N-acetylhexosamine phosphatidyl hexose
PHexHex2NAc: di-N-acetylhexosamine phosphatidyl hexose
PHexHex+C5H8: glycosylated phosphatidyl hexose bearing an additional mass of 68
CDP: cytidine diphosphate
PG: phosphatidyl glycerol
C_21_-PC: phosphatidyl choline diacylglycerol with two C_21_ fatty acid chains
B&D: Bligh and Dyer
MeOH: methanol
DCM: dichloromethane
ACN: acetonitrile
TFA: trifluoroacetic acid
FA: formic acid
NH_3_: ammonium hydroxide or aqueous ammonia
DHB: 2,3-dihydroxybenzoic acid
UHPLC: ultra-high performance liquid chromatography
Q-TOF: quadrupole time of flight
QQQ: triple quadrupole
MS: mass spectrometry
ESI: electrospray ionization
APCI: atmospheric-pressure chemical ionization
MALDI: matrix-assisted laser desorption/ionization
FT: Fourier transform
ICR: ion cyclotron resonance.

