## Supplementary file for "The exploration of *Thermococcus barophilus* lipidome reveals the widest variety of phosphoglycolipids in Thermococcales"

### *Supplementary material*

#### Supplementary text

Numerous mass-to-charge ratios ranging from *m/z* = 1786.4 to 1800.4 with 1 mass unit increments were detected in the UHPLC-MS unresolved retention time window (23 to 27 min). The major component of this unresolved peak was compound XIX (PI-GDGT0-PI, $[M+H]^+$ at *m/z* = 1786.367 and $[M+NH_4]^+$ at *m/z* = 1803.382; Figure S9). No fragmentation pattern was available for the peaks of higher masses, but two non mutually exclusive hypotheses might be suggested for the structures of these slightly heavier tetraether-based IPLs. On the one hand, archaeal tetraether-based IPL are composed of at least 86 carbon atoms (in the case of core GDGT) and are thus highly likely to present one or more $^{13}C$ isotopologues, which masses differ by one or more units compared to the $^{12}C$ monoisotopic form. The mass distribution in the PI-GDGT-PI peak indeed matched a theoretical isotopic pattern for this compound (Figure S10), supporting the presence of isotopologues with up to six $^{13}C$. On the other hand, *T. barophilus* and other archaea were shown to produce diverse tetraether core structures, notably GTGT (28). In particular, GTGT0 molecular mass is shifted upwards by two mass units relative to GDGT0 (1304.34 vs. 1302.32). The hydrolysis experiments carried out on the total lipid extract and the biomass of *T. barophilus* systematically yielded GTGT0 XXII but no ring-containing tetraethers (Figure 4; 24), supporting its ability to synthesize GTGT0-based IPLs. We thus suggested compound XXIII to be PI-GTGT0-PI, with a molecular mass of *m/z* = 1788.376. The remaining molecular masses, i.e., 1787.4, 1789.4 and 1790.4 and 1804.4, 1806.4 and 1807.4 would consequently correspond to protonated and ammoniated adducts of $^{13}C$-containing PI-GDGT0-PI XIX and PI-GTGT0-PI XIII.

#### Supplementary figures

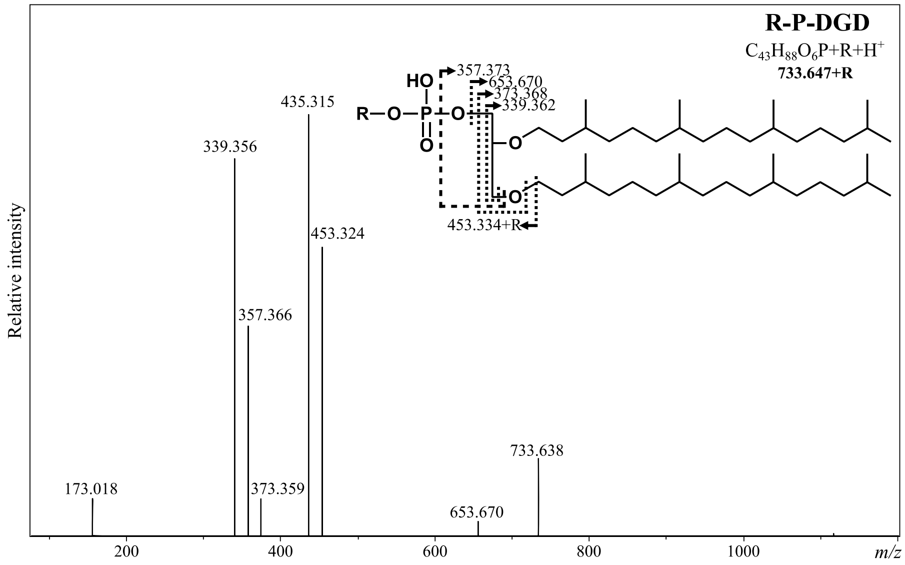


**Figure S1: Diagnostic molecular ions of diether-based phospholipids.**
IPLs were separated using UHPLC and were detected using ESI-MS operated in positive mode. $MS^2$ scans were automatically generated by fragmentation of the most abundant ions at each scan. A representative mass spectrum is displayed. Insert indicates the compound structure with fragmentation, chemical formula, and total molecular ion mass. Characteristic ions found in every diether-based phospholipids are displayed.

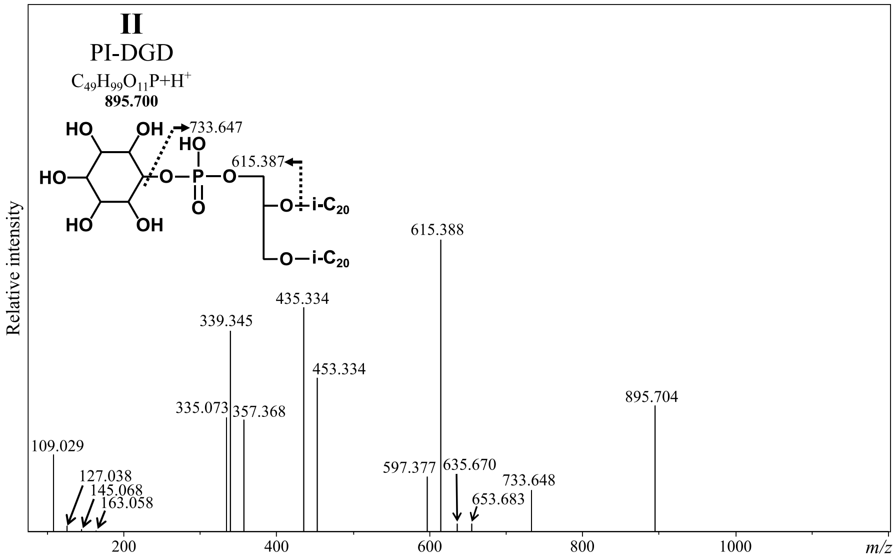
II
PI-DGD
C49H99O11P+H+
895.700
733.647
615.387
i-C20
Relative intensity
615.388
435.334
339.345
453.334
895.704
335.073
357.368
109.029
597.377
635.670
733.648
127.038
145.068
163.058
653.683
200
400
600
800
1000
m/z

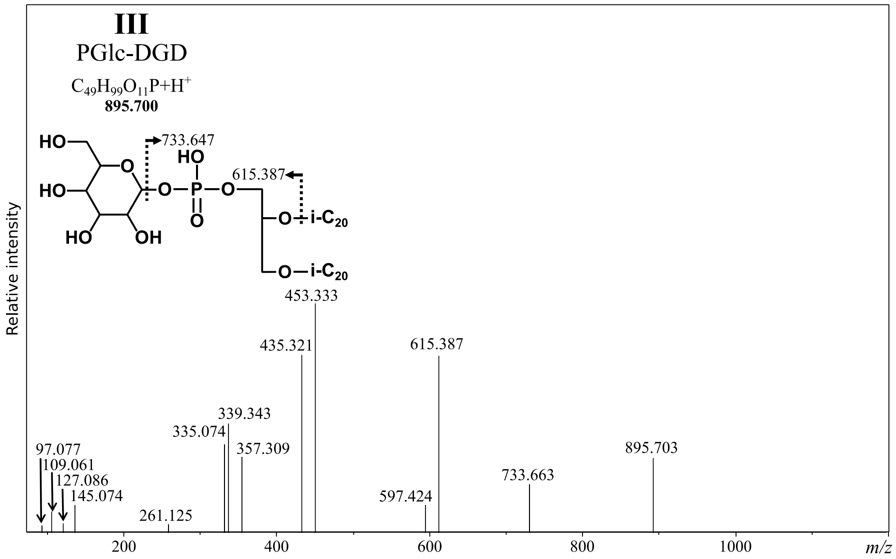
III
PGlc-DGD
C49H99O11P+H+
895.700
733.647
615.387
i-C20
Relative intensity
453.333
435.321
615.387
339.343
335.074
357.309
895.703
97.077
109.061
127.086
145.074
733.663
597.424
261.125
200
400
600
800
1000
m/z

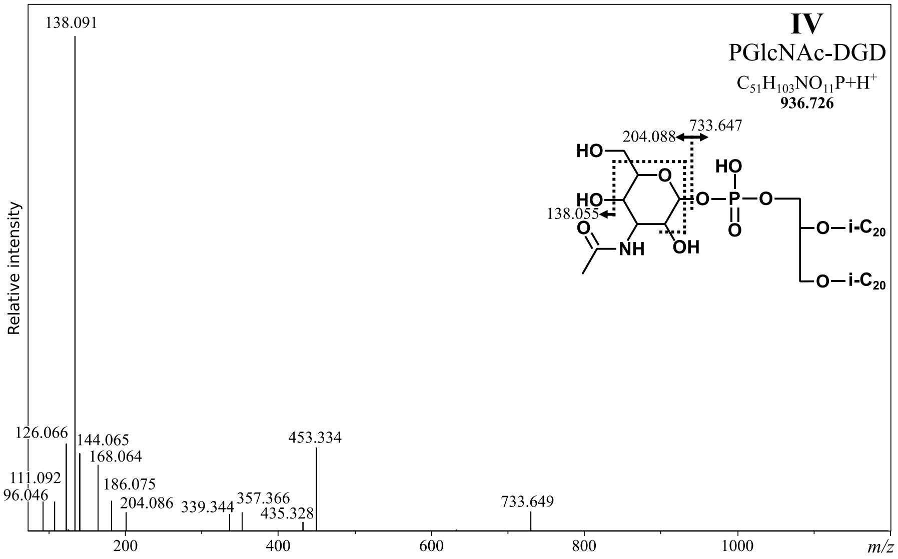
IV
PGlcNAc-DGD
C51H103NO11P+H+
936.726
204.088
733.647
138.055
i-C20
Relative intensity
138.091
126.066
144.065
168.064
453.334
111.092
96.046
186.075
204.086
339.344
357.366
435.328
733.649
200
400
600
800
1000
m/z

**Figure S2: Diagnostic molecular ions of PI-DGD II, PGlc-DGD III and PGlcNAc-DGD IV.**

IPLs were separated using UHPLC and were detected using ESI-MS operated in positive mode. $MS^2$ scans were automatically generated by fragmentation of the most abundant ions at each scan. A representative mass spectrum is displayed for each compound. Inserts indicate the compound structure with fragmentation, chemical formula, and total molecular ion mass. Fragments already present in previous figures are not indicated although their masses were detected (refer to Figures S1 and S2). Note the similarity between PI-DGD II and PGlc-DGD III and the differences in the polar headgroup characteristic ions of PGlcNAc-DGD IV. The N-acetylation could be located on the sugar residue of PGlcNAc-DGD IV, but its exact location could not be properly assessed and was thus arbitrarily drawn.

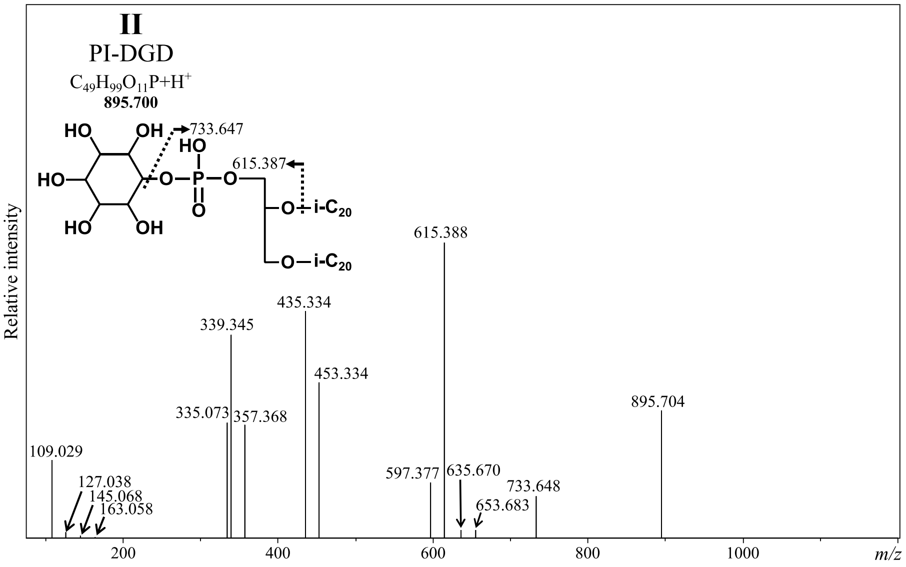

II
PI-DGD
C49H99O11P+H+
895.700
733.647
615.387
i-C20
i-C20
Relative intensity
615.388
435.334
339.345
453.334
335.073
357.368
895.704
109.029
127.038
145.068
163.058
597.377
635.670
653.683
733.648
200
400
600
800
1000
m/z


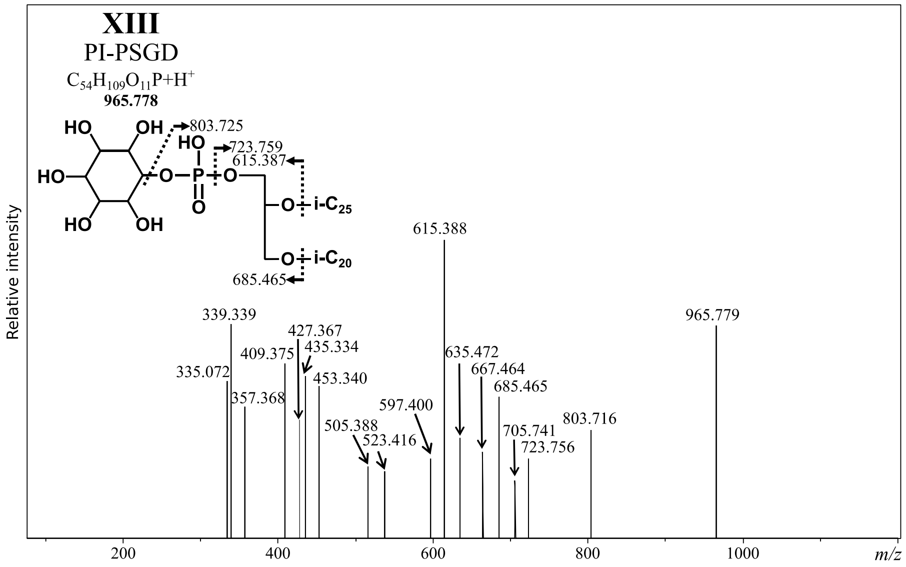

XIII
PI-PSGD
C54H109O11P+H+
965.778
803.725
723.759
615.387
i-C25
i-C20
685.465
Relative intensity
615.388
965.779
339.339
427.367
435.334
409.375
335.072
453.340
357.368
635.472
667.464
685.465
597.400
505.388
523.416
803.716
705.741
723.756
200
400
600
800
1000
m/z


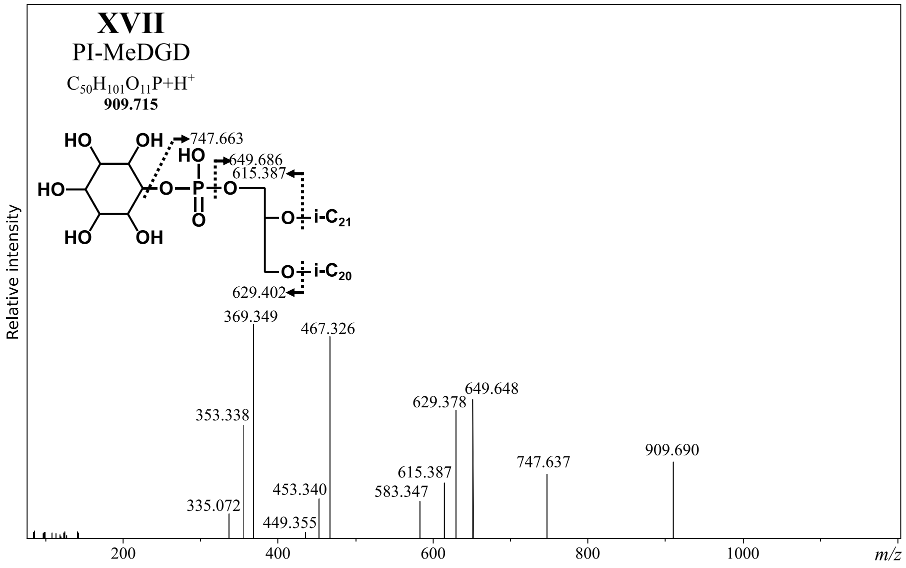

XVII
PI-MeDGD
C50H101O11P+H+
909.715
747.663
649.686
615.387
i-C21
i-C20
629.402
Relative intensity
369.349
467.326
649.648
629.378
353.338
909.690
747.637
615.387
453.340
583.347
335.072
449.355
200
400
600
800
1000
m/z

**Figure S3: Diagnostic molecular ions of PI-DGD II, PI-PSGD XIII and PI-MeDGD XVII.**

IPLs were separated using UHPLC and were detected using ESI-MS operated in positive mode. $MS^2$ scans were automatically generated by fragmentation of the most abundant ions at each scan. A representative mass spectrum is displayed for each compound. Inserts indicate the compound structure with fragmentation, chemical formula, and total molecular ion mass. Note the differences in the core structure characteristic ions. Fragments already present in previous figures are not indicated although their masses were detected (refer to Figure S1). The configuration of the $C_{20}$ and $C_{25}$ isoprenoid chains on the glycerol backbone of PI-PSGD XIII could not be properly assessed and was thus arbitrarily drawn. The position of the additional methylation and the configuration of the $C_{20}$ and $C_{21}$ isoprenoid chains on the glycerol backbone of PI-MeDGD XVII could not be properly assessed and was thus arbitrarily drawn.

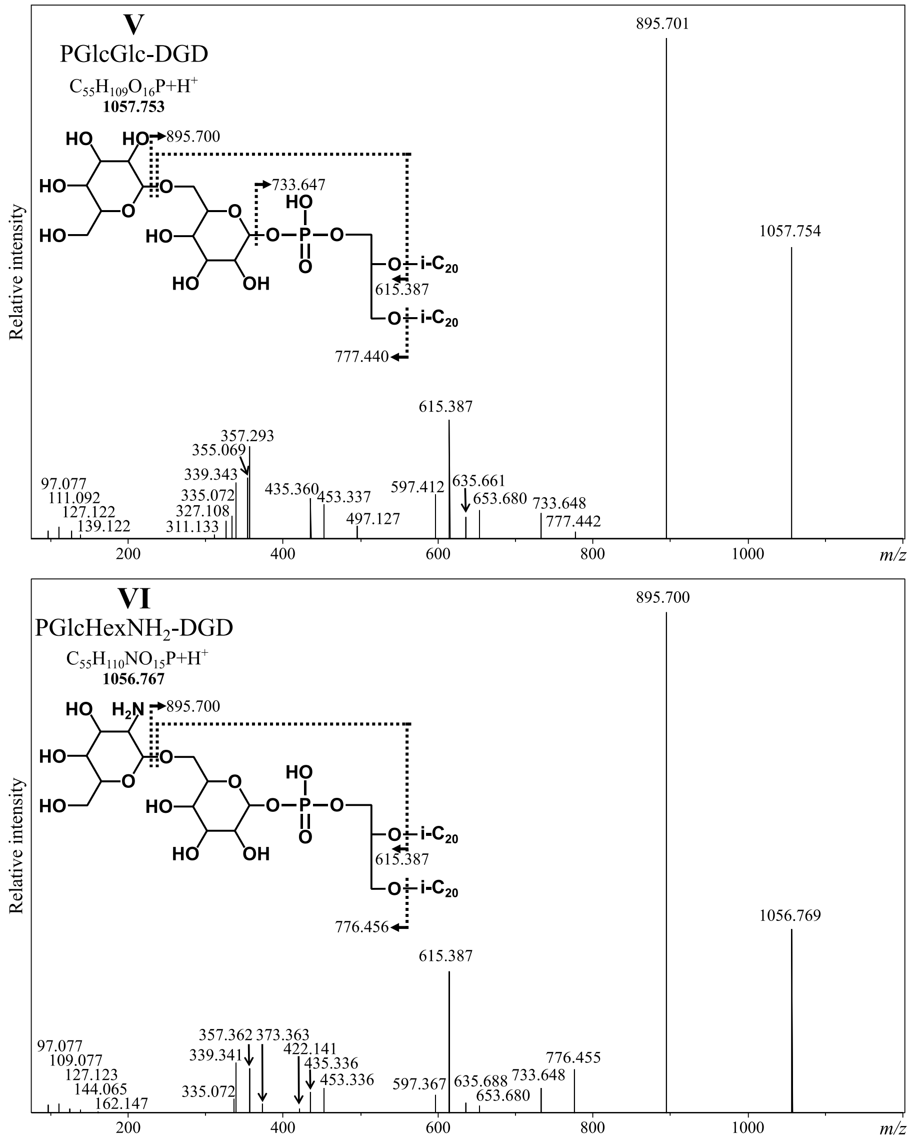


**Figure S4: Diagnostic molecular ions of PGlcGlc-DGD V and PGlcHexNH$_2$-DGD VI,.**

IPLs were separated using UHPLC and were detected using ESI-MS operated in positive mode. MS$^2$ scans were automatically generated by fragmentation of the most abundant ions at each scan. A representative mass spectrum is displayed for each compound. Inserts indicate the compound structure with fragmentation, chemical formula, and total molecular ion mass. Fragments already present in previous figures are not indicated although their masses were detected (refer to Figures S1 and S3). The NH$_2$ group of PGlcHexNH$_2$-DGD VI could be located on the second hexose moiety, but its exact location could not be properly assessed and was thus arbitrarily drawn.

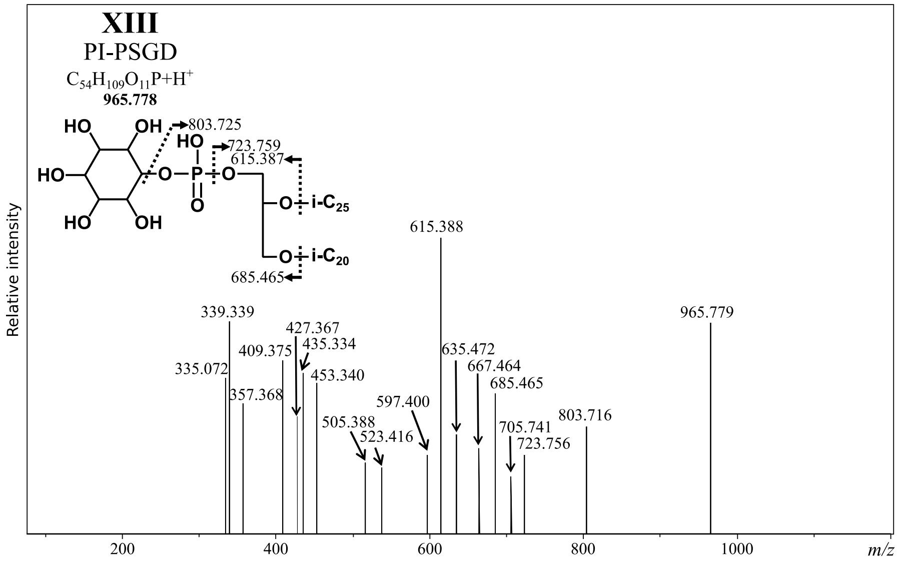
XIII
PI-PSGD
$C_{54}H_{109}O_{11}P+H^+$
965.778
Relative intensity
m/z

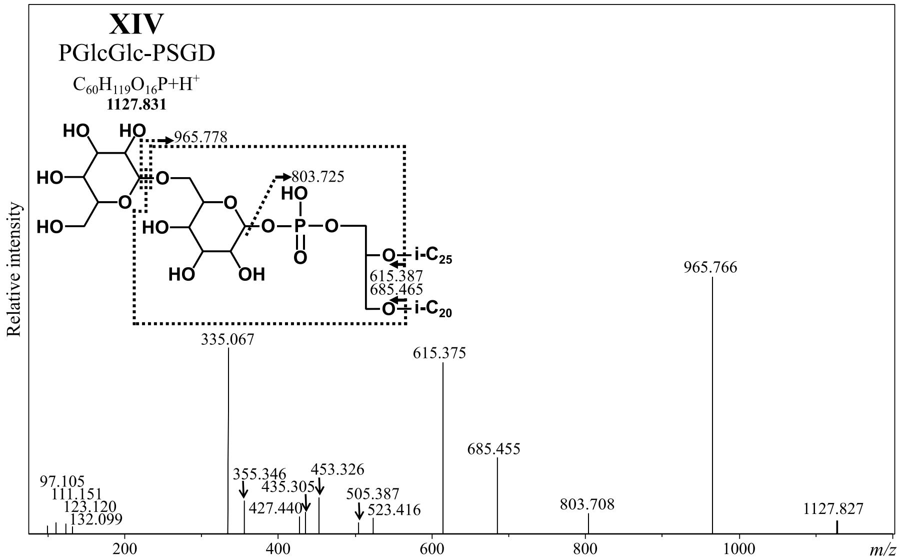
XIV
PGlcGlc-PSGD
$C_{60}H_{119}O_{16}P+H^+$
1127.831
Relative intensity
m/z

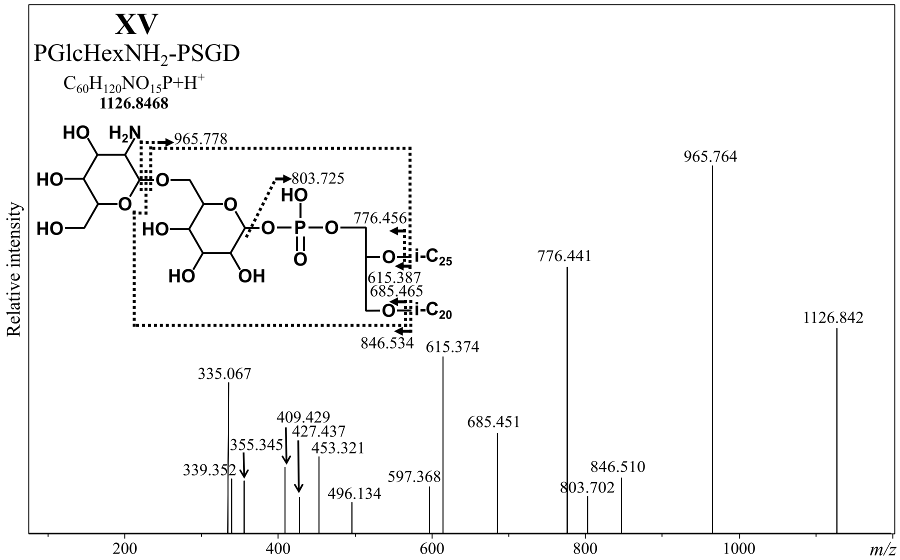
XV
PGlcHexNH2-PSGD
$C_{60}H_{120}NO_{15}P+H^+$
1126.8468
Relative intensity
m/z

**Figure S5: Diagnostic molecular ions of PI-PSGD XIII, PGlcGlc-PSGD XIV and PGlcHexNH$_2$-PSGD XV.**

IPLs were separated using UHPLC and were detected using ESI-MS operated in positive mode. MS$^2$ scans were automatically generated by fragmentation of the most abundant ions at each scan. A representative mass spectrum is displayed for each compound. Inserts indicate the compound structure with fragmentation, chemical formula, and total molecular ion mass. Fragments already present in previous figures are not indicated although their masses were detected (refer to Figures S1, S2, and S4). Note the presence of ions at 615.387 and 685.465 indicative of a PSGD core lipid for the three compounds. The NH$_2$ group of PGlcHexNH$_2$-PSGD XV could be located on the second hexose moiety, but its exact location could not be properly assessed and was thus arbitrarily drawn.

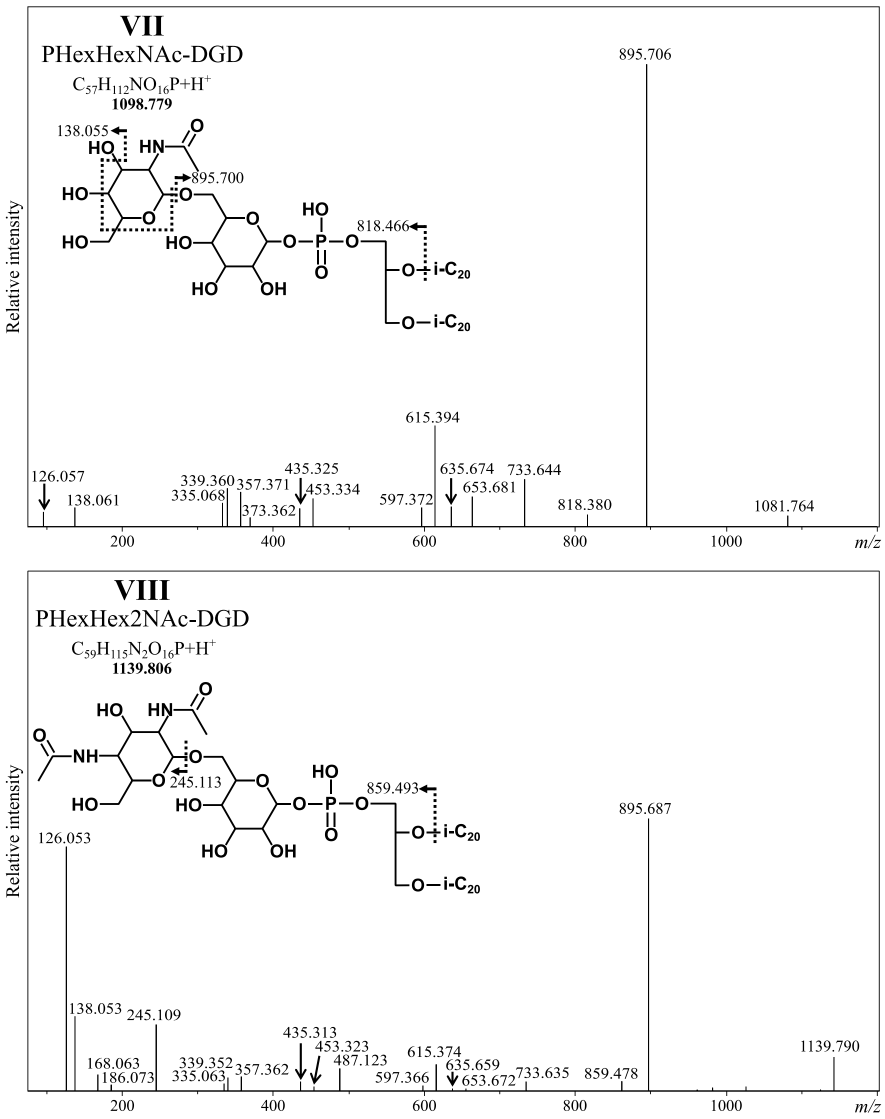


**Figure S6: Diagnostic molecular ions of PHexHexNAc-DGD VII and PHexHex2NAc-DGD VIII.** IPLs were separated using UHPLC and were detected using ESI-MS operated in positive mode. $MS^2$ scans were automatically generated by fragmentation of the most abundant ions at each scan. A representative mass spectrum is displayed for each compound. Inserts indicate the compound's structure with fragmentation, chemical formula, and total molecular ion mass. Fragments already present in previous figures are not indicated although their masses were detected (refer to Figures S1 and S4). The N-acetylations could be located on the second hexose moiety, but their exact location could not be properly assessed and was thus arbitrarily drawn.

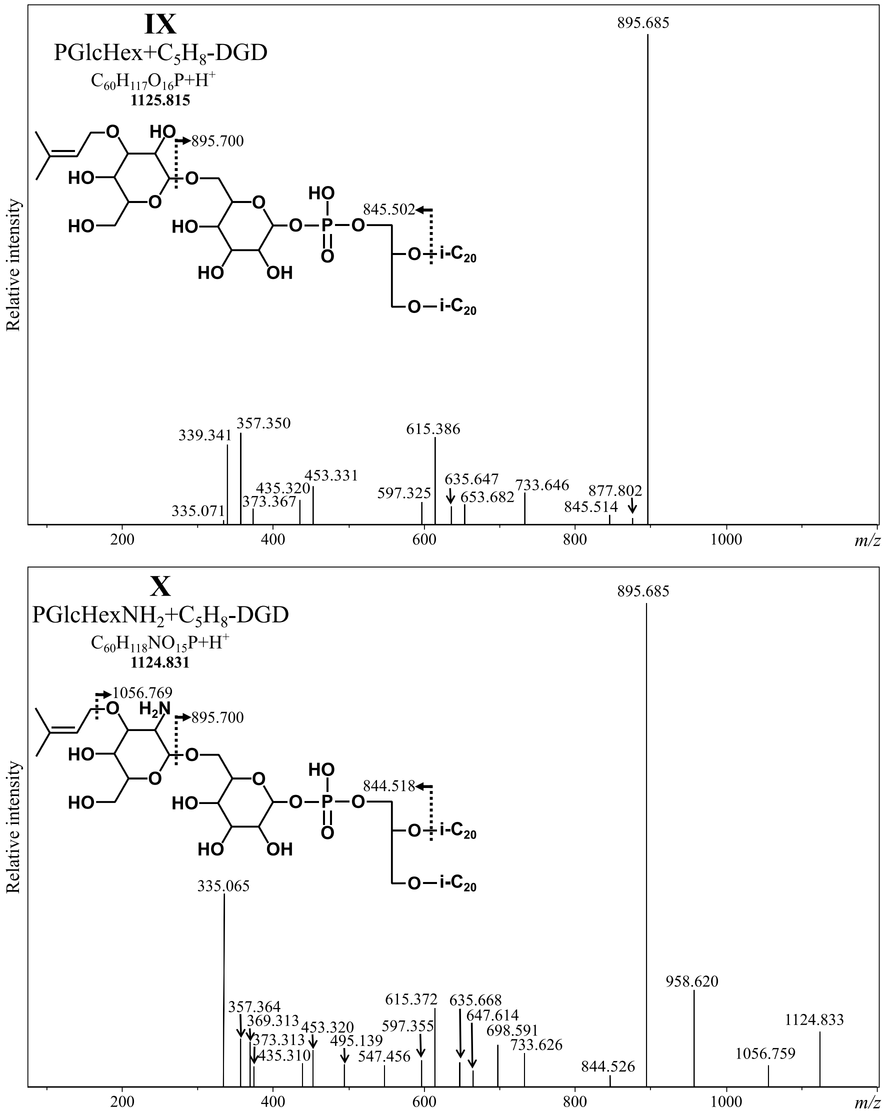


**Figure S7: Diagnostic molecular ions of PGlcHex+$C_5H_8$-DGD IX and PHexHexNH$_2$+$C_5H_8$ X.**

IPLs were separated using UHPLC and were detected using ESI-MS operated in positive mode. $MS^2$ scans were automatically generated by fragmentation of the most abundant ions at each scan. A representative mass spectrum is displayed for each compound. Inserts indicate the compound structure with fragmentation, chemical formula, and total molecular ion mass. Fragments already present in previous figures are not indicated although their masses were detected (Figures S1 and S4). Compounds IX and X showed the same masses than PGlcGlc-DGD V and PGlcHexNH$_2$-DGD **VI** shifted upwards by 68 mass units, respectively (Figure S4). The nature of these 68 additionnal mass units could not be ascertained but was assigned to an isoprene group. The isoprene groups could be located on the second hexose moiety, but their exact location could not be properly assessed and was thus arbitrarily drawn.

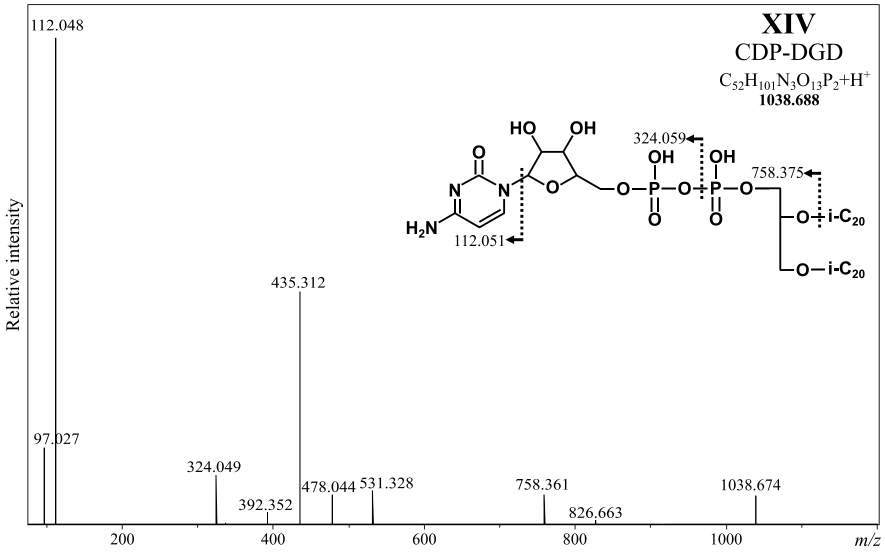


**Figure S8: Diagnostic molecular ions of CDP-DGD XI.**

IPLs were separated using UHPLC and were detected using ESI-MS operated in positive mode. $MS^2$ scans were automatically generated by fragmentation of the most abundant ions at each scan. A representative mass spectrum is displayed. Inserts indicate the compound's structure with fragmentation, chemical formula, and total molecular ion mass. Fragments already present in previous figures are not indicated although their masses were detected (Figure S1).

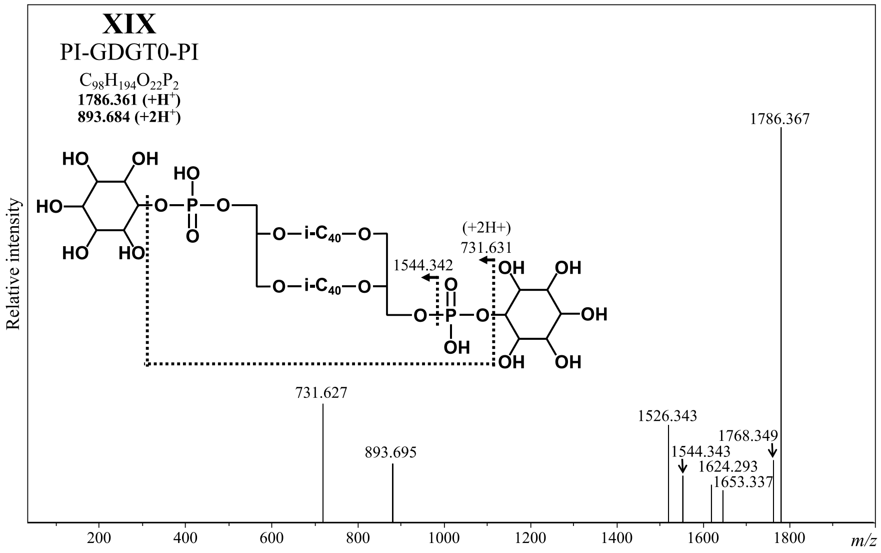


**Figure S9: Diagnostic molecular ions of PI-GDGT0-PI XIX.**

IPLs were separated using UHPLC and were detected using ESI-MS operated in positive mode. $MS^2$ scans were automatically generated by fragmentation of the most abundant ions at each scan. A representative mass spectrum is displayed. Inserts indicate the compound structure with fragmentation, chemical formula, and total molecular ion mass.

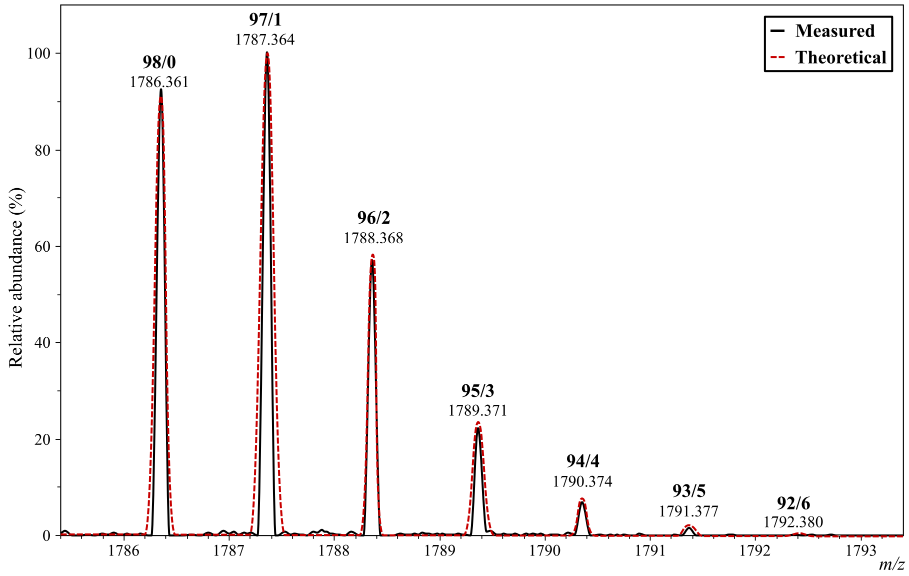


**Figure S10: Isotopolog pattern of PI-GDGT0-PI XIX in *Thermococcus barophilus*.**

Comparison of the averaged mass distribution in the PI-GDGT-PI XIX peak (black line) with a theoretical isotopic pattern (red dashed line) obtained using the IsotopePattern built-in function of the DataAnalysis software (chemical formula, $C_{98}H_{194}O_{22}P_2$; $[M+H]^+$ mode; isotope threshold, 0.1 %). Detected $[M+H]^+$ and numbers of $^{12}C/^{13}C$ are indicated above the corresponding peaks. The monoisotopic ion mass of PI-GDGT-PI XIX is 1786.361 (refer to Figure 1).

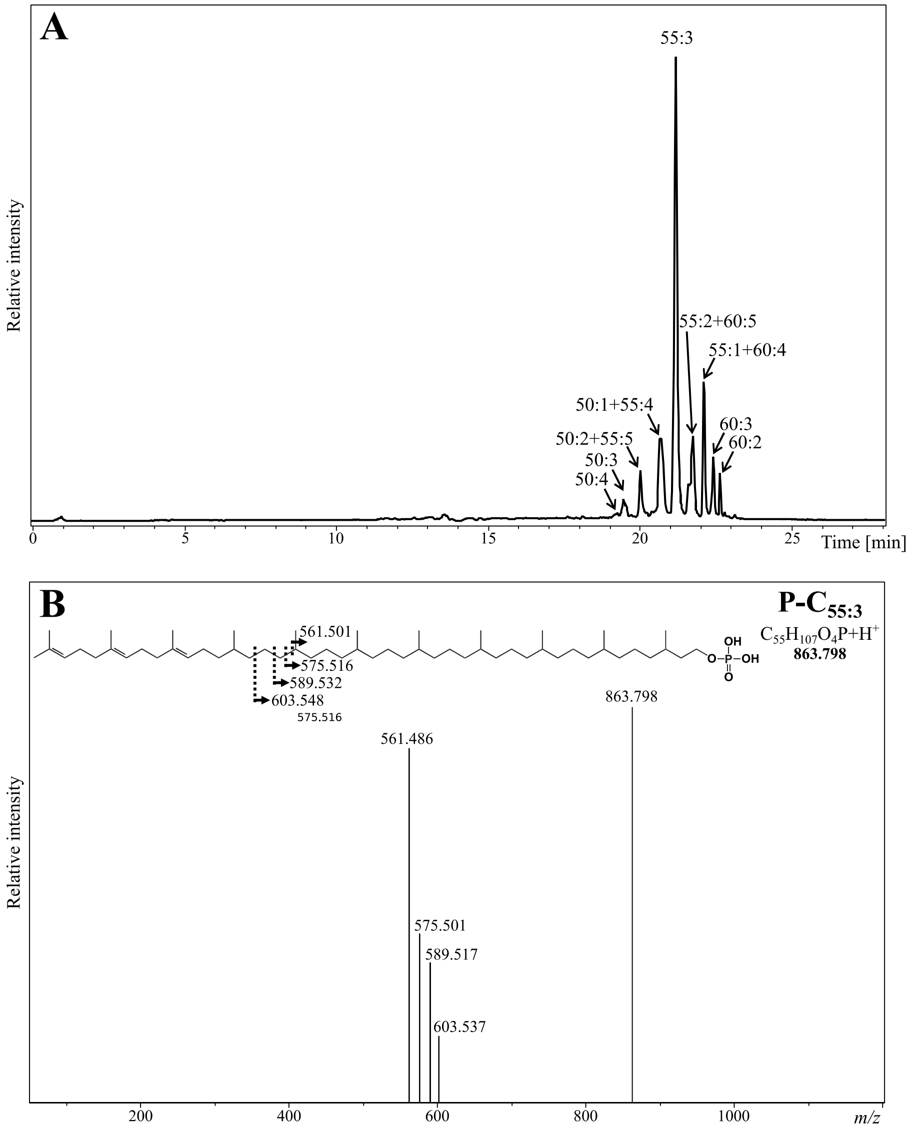


**Figure S11: *Thermococcus barophilus* synthesizes polyprenyl monophosphates containing 10 to 12 isoprene units and various unsaturation degrees.**

*T. barophilus*' polyprenyl derivatives were extracted alongside IPLs with the same extraction procedure but were separated using reverse phase UHPLC with mixtures of MeOH:$H_2O$:FA:$NH_3$ (85:15:0.04:0.1, *v*/*v*/*v*/*v*) and propan-2-ol:MeOH:FA:$NH_3$ (50:50:0.04:0.1, *v*/*v*/*v*/*v*; for further details refer to the Methods section). (**A**) The UHPLC chromatogram was drawn in positive mode by extracting the following protonated ions with a mass deviation of 0.02 Da: 791.70, 793.72, 795.74, 797.75, 859.77, 861.78, 863.80, 865.81, 867.83, 929.84, 931.86, 933.88, and 935.89. (**B**) Diagnostic molecular ions of the most abundant polyprenyl of *T. barophilus*, i.e., with a phosphatidyl head group, 11 isoprene units, and three unsaturations (P-$C_{55:3}$). Insert indicates the compound structure with fragmentations, chemical formulas, and total molecular ion masses.

#### Supplementary tables

**Table S1: Triple quadrupole parameters and detection windows selected for each transition of the sMRM method. DP: declustering potential, EP: entrance potential, CE: collision energy, CXP: collision cell exit potential.**

| Carbohydrate | | Abbreviation | Transition for quantification | DP | EP | CE | CXP | Dwell weight | Detection window (min) |
|---|---|---|---|---|---|---|---|---|---|
| Hexoses (M = 180.2 g.mol$^{-1}$) | Glucose | Glc | 179.1 -> 89.0 | -20 | 10 | -10 | -10 | 1.5 | 9.0 – 21 |
| | Galactose | Gal | | | | | | | |
| | Gulose | Gul | | | | | | | |
| | Tagatose | Tag | | | | | | | |
| | Fructose | Fru/Alp | | | | | | | |
| | Allose | | | | | | | | |
| | Mannose | Man | 179.1 -> 119.0 | -10 | 10 | -10 | -10 | 1.5 | 9.0 – 21 |
| Pentoses (M = 150.1 g.mol$^{-1}$) | Lyxose | Lyx | 149.0 -> 58.9 | -15 | 10 | -20 | -5 | 1 | 7.0 – 11 |
| | Xylose | Xyl/Ara | | | | | | | |
| | Arabinose | | | | | | | | |
| Deoxyhexose (M = 164.2 g.mol$^{-1}$) | Rhamnose | Rha | 163.1 -> 59.0 | -10 | 10 | -20 | -10 | 1 | 4.5 – 7.5 |
| Sugar alcohol (M = 180.2 g.mol$^{-1}$) | Inositol | Ino | 179.0 -> 86.8 | -58 | 10 | -23 | -7 | 1 | 29.3 – 33.3 |
| Disacchararide (M = 342.3 g.mol$^{-1}$) | Saccharose | Sac | 341.1 -> 59.0 | -30 | 10 | -55 | -5 | 1 | 30.0 – 32.0 |
| Acetylated aminosugar (M = 221.2 g.mol$^{-1}$) | N-acetylglucosamine | GlcNAc | 220.1 -> 118.6 | -10 | 10 | -10 | -11 | 1 | 13.5 – 17.5 |
